# Single-cell Networks Reorganise to Facilitate Whole-brain Supercritical Dynamics During Epileptic Seizures

**DOI:** 10.1101/2021.10.14.464473

**Authors:** DRW Burrows, G Diana, B Pimpel, F Moeller, MP Richardson, DS Bassett, MP Meyer, RE Rosch

## Abstract

Excitation-inhibition (EI) balance may be required for the organisation of brain dynamics to a phase transition, criticality, which confers computational benefits. Brain pathology associated with EI imbalance may therefore occur due to a deviation from criticality. However, evidence linking critical dynamics with EI imbalance-induced pathology is lacking. Here, we studied the effect of EI imbalance-induced epileptic seizures on brain dynamics, using *in vivo* whole-brain 2-photon imaging of GCaMP6s larval zebrafish at single-neuron resolution. We demonstrate the importance of EI balance for criticality, with EI imbalance causing a loss of whole-brain critical statistics. Using network models we show that a reorganisation of network topology drives this loss of criticality. Seizure dynamics match theoretical predictions for networks driven away from a phase transition into disorder, with the emergence of chaos and a loss of network-mediated separation, dynamic range and metastability. These results demonstrate that EI imbalance drives a pathological deviation from criticality.

## Introduction

A delicate balance between excitation and inhibition supports the diverse repertoire of neural dynamics needed for behaviour. This excitation-inhibition (EI) balance is thought to be important for healthy brain function as multiple neurodevelopmental disorders are linked to EI imbalance, including schizophrenia, autism and epilepsy (Fritschy, 2008; Gao & Penzes, 2015; Lee et al., 2017; Žiburkus et al., 2013). EI balance manifests synaptically as invariant ratios of excitatory-inhibitory synapses along dendritic segments (Iascone et al., 2020). Interestingly, synaptic EI balance can shape global dynamics, as demonstrated by the emergence of generalised seizures in synaptopathies (Escayg & Goldin, 2010; Johannesen et al., 2016; R. Rosch et al., 2019). At present, our understanding of the role of EI balance in shaping dynamics and ultimately computation comes from small population recordings, suggesting roles in shaping receptive fields (B. Liu et al., 2010), supporting coincidence detection (Wehr & Zador, 2003) and enabling gain control (Bhatia et al., 2019). How synaptic EI balance shapes population dynamics in whole-brain networks to facilitate computation remains an open question, due to the technical difficulties of collecting and analysing such high dimensional datasets.

One appealing approach is to borrow concepts from statistical physics, which measures microscopic variables probabilistically to estimate macroscale properties of complex systems. Using such approaches, it has been claimed that neuronal populations exhibit dynamics analogous to atomic spins in a ferromagnetic lattice organised to a phase transition between order and chaos, known as criticality (Bak et al., 1987, 1988; Hesse & Gross, 2014; Sethna et al., 2001). In fact, various statistical signatures of criticality have been documented empirically in neural recordings across diverse scales (Beggs & Plenz, 2003; Kitzbichler et al., 2009; Meisel et al., 2012; Ponce-Alvarez et al., 2018). However, such statistical indicators of criticality are purely correlative and therefore the extent to which brain dynamics share universal properties of idealised, physical systems operating exactly at a phase transition is unclear (Fontenele et al., 2019; Wilting & Priesemann, 2019). Nonetheless, a system operating within a collection of near-to-critical states offers a biologically plausible regime of brain activity which can support computation (Gross, 2021; Kinouchi et al., 2020; Moretti & Muñoz, 2013; Priesemann et al., 2014). Interestingly, *in silico* evidence suggests that only EI balanced networks support scale-invariant avalanches (Poil et al., 2012), spatio-temporal cascades of neural activity that span the entire scale of the system, a defining feature of criticality. Furthermore, perturbing EI balance *in vitro* removes signatures of criticality (Haldeman & Beggs, 2005; Shew et al., 2011). Therefore, the tendency of the brain to homeostatically maintain EI balance (Turrigiano & Nelson, 2004) could serve to tune its dynamics near to a phase transition. The role of EI balance in shaping near-critical dynamics has yet to be tested *in vivo*.

Systems at criticality maximise various computational capacities, such as: i) network-mediated separation, the ability to separate similar inputs to distinguishable outputs (Bertschinger & Natschläger, 2004; Maass et al., 2002), ii) dynamic range, the range of inputs that the network can encode (Kinouchi & Copelli, 2006; Shew et al., 2011), and iii) metastability, the tendency of the brain to transiently explore a diversity of semi-stable states supporting flexible dynamics (Deco et al., 2017; Fingelkurts & Fingelkurts, 2001). Therefore, it has been theorised that EI imbalance-associated brain dysfunction may emerge as a deviation from the critical state (Zimmern, 2020). In fact, epileptic seizures, which can emerge due to pathological EI imbalances, are associated with a loss of critical statistics in EEG and MEG recordings (Arviv et al., 2016; Meisel et al., 2012). This has led to the prediction that seizure initiation manifests as a supercritical state (Beggs & Plenz, 2003), where dynamics move away from a transition point into chaos (Haldeman & Beggs, 2005), thus causing inputs to exponentially grow in time and saturate the system (Harris, 1963). Whilst some features of supercritical dynamics have been reported using macroscale seizure recordings, such techniques coarse-grain the underlying dynamics which obscures the heterogeneity of cellular activity and could alter critical statistics (Keller et al., 2010; Meyer et al., 2018; Muldoon et al., 2013). Conversely, recording from subregions of a full system can produce spurious critical statistics (Priesemann et al., 2009), while distinct brain regions may reside at different near-critical states (Suryadi et al., 2018). Therefore, to make strong claims about the critical nature of entire neuronal systems, we require single cell recordings with whole-brain coverage. Whether single neuron dynamics collectively engage to give rise to a supercritical state in whole-brain networks during seizures is untested. Showing this experimentally, would demonstrate that synaptic EI imbalance can impair computation by disrupting near-critical dynamics across the whole brain.

Here we take advantage of the transparency of the larval zebrafish to perform *in vivo* functional imaging of the whole brain at single cell resolution (Ahrens et al., 2013). We test 2 key hypotheses: i) EI balance regulates near-critical dynamics, and ii) epileptic seizures emerge as a supercritical state which impairs brain computation. We find that perturbations to EI balance give rise to abrupt changes in critical statistics, suggesting a role of EI balance in organising dynamics near to a phase transition. Furthermore, epileptic seizures manifest as a chaotic state giving rise to reduced network-mediated separation, dynamic range and metastability, demonstrating that seizures emerge as a supercritical state.

## Results

To study critical dynamics *in vivo* we analysed neuronal activity of GCaMP6s-expressing larval zebrafish captured via whole-brain 2-photon imaging at single cell resolution (See Figure S1). To perturb EI balance we used the GABA_A_ receptor antagonist pentylenetetrazole (PTZ), which causes epileptiform discharges (Baraban et al., 2005; R. E. Rosch et al., 2018). We recorded 3 x 30 minute consecutive imaging blocks: 1) *spontaneous* activity representing EI balance, 2) *5mM PTZ* causing EI imbalance giving rise to focal seizures, and 3) *20mM PTZ* causing EI imbalance giving rise to generalised seizures (See Figure S2) (Diaz Verdugo et al., 2019). We segmented ~9,000 neurons per dataset, estimating their latent on/off states using a hidden Markov model (HMM) (Diana et al., 2019) (See Figure S1D & 1B). This allowed us to measure the propagation of neuronal avalanches and the evolution of population dynamics during EI balance and imbalance (See Methods & Figure 1).

**F1.**
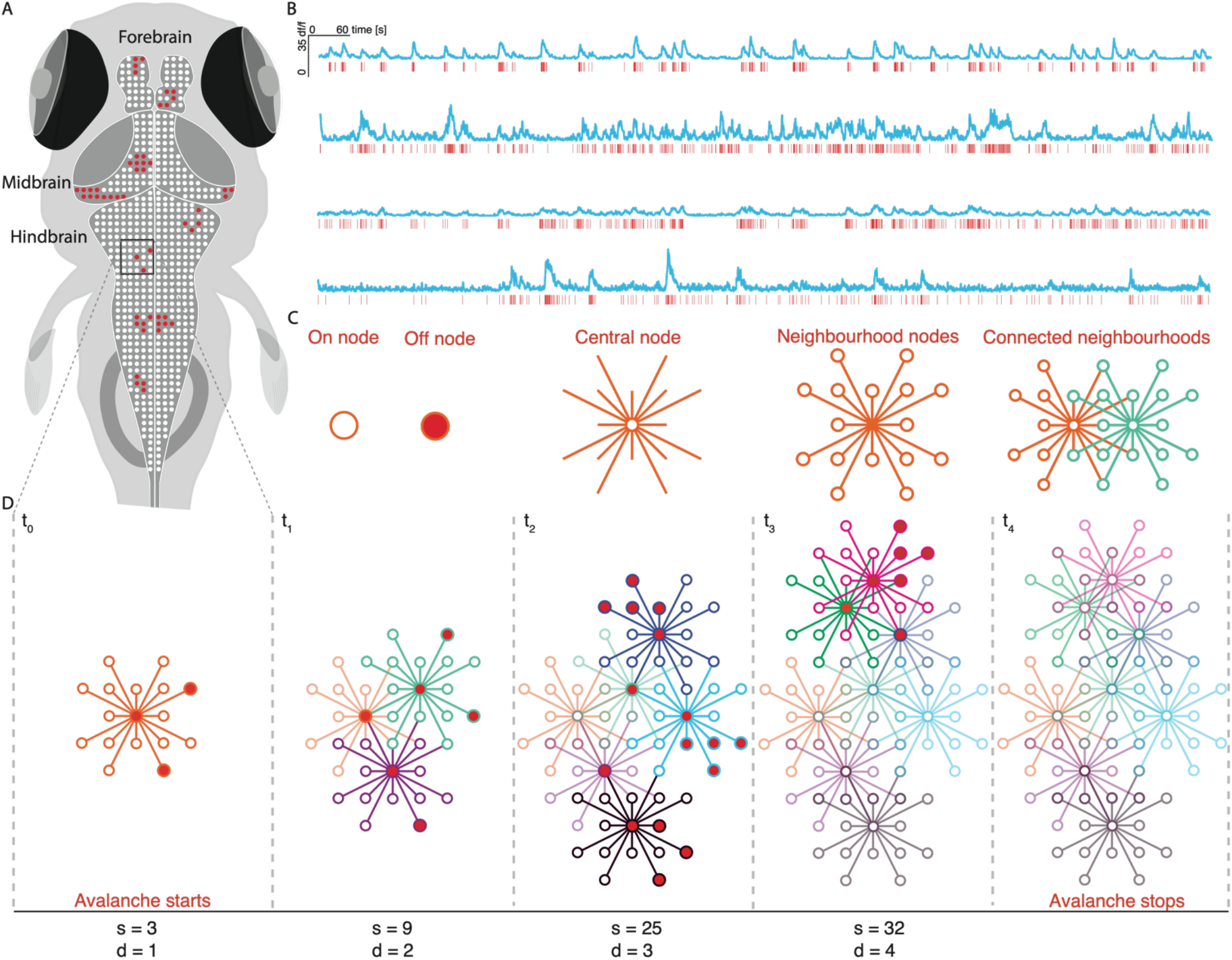
Avalanche estimation. (A) Neurons exhibit on (red circles) and off (white circles) dynamics giving rise to ensembles that grow in space and time. (B) HMM estimation of calcium transients showing normalised traces (blue), and estimated calcium transients (red). (C) Avalanche calculation legend. (D) For avalanches to begin, at least 3 nodes within the same neighbourhood must be active at *t_0_*. To propagate in time, any avalanche node active at *t_0_* must also be active at *t_1_*. Once this step is satisfied, any nodes active at *t_1_* that are connected via a neighbourhood to avalanche nodes at *t_1_* become avalanche members. Avalanches terminate when no more nodes are active. Avalanche size (s) is the total number of activations and duration (d) is the number of time steps of the avalanche.

### Spontaneous Neural Dynamics Exhibit Critical Statistics

Several statistical features have been reported as indicators of criticality, including (i) power-law probability distributions of avalanche size and duration, (ii) scaling relationships between avalanche power-law exponents, (iii) branching ratios close to unity and (iv) power-law scaling of neuron correlation and distance. We evaluate each of these features to validate the presence of near-critical dynamics in spontaneous activity.

A key feature of criticality is the presence of scale-invariant neuronal avalanches, contiguous clusters of activity which propagate in space and time (See Figure 1D), giving rise to power-law probability distributions. We estimated probability distributions for avalanche size and duration from calcium imaging data, which were well fit by power-laws (See Figure 2A). Using log likelihood ratio testing, we found that all datasets were better explained by power-law than lognormal distributions, the most rigorous alternative heavy-tail distribution test (See Methods) (Alstott et al., 2014). Importantly, measuring neuronal avalanches from sequences of oscillatory peaks in human intracranial recordings also reveals power-law statistics in baseline activity (See Figure S3B). This indicates the robustness of power-law relationships in neural activity across brain scales.

**F2.**
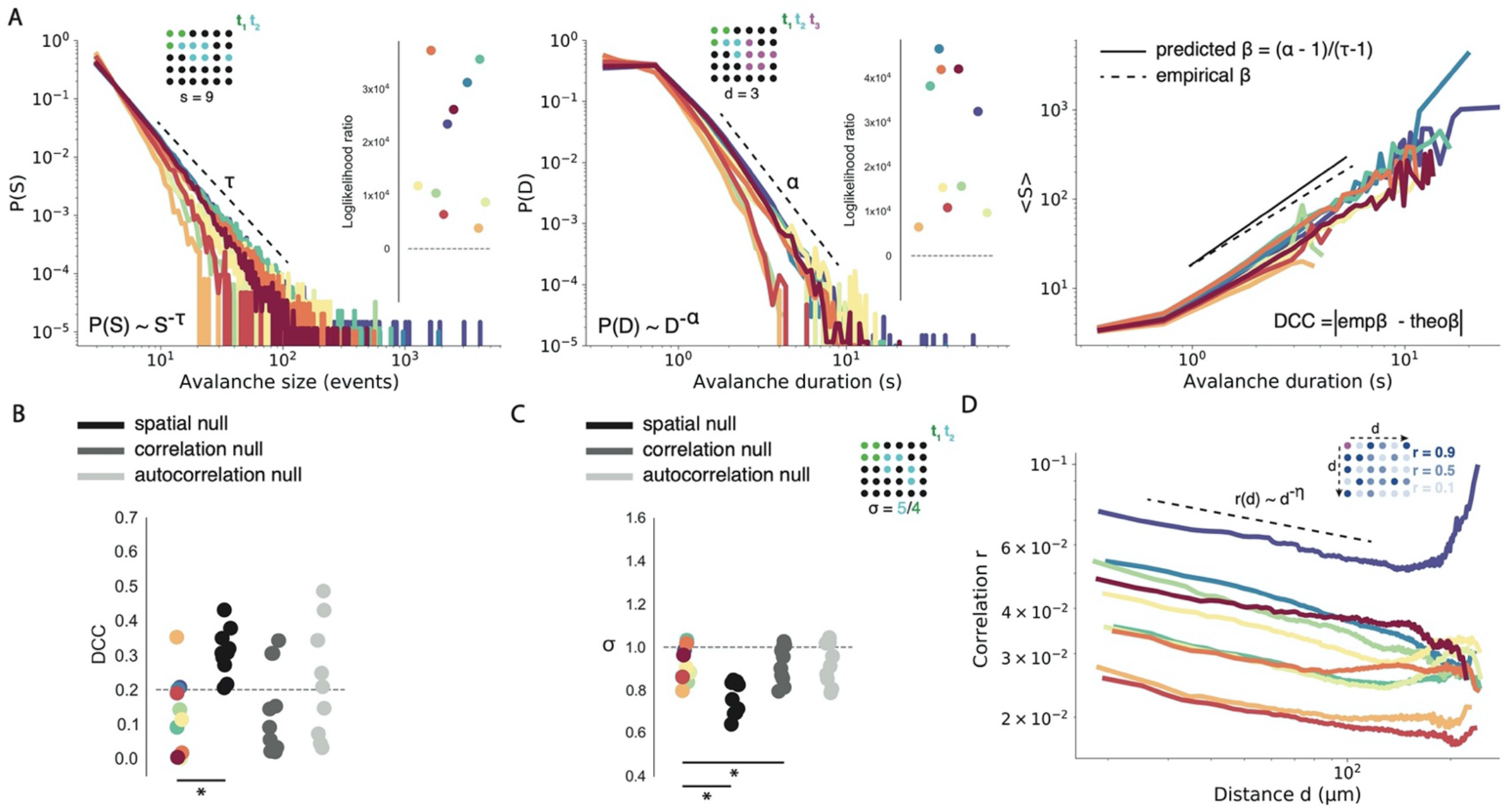
Spontaneous dynamics exhibit critical statistics. (A) Probability distributions for avalanche size (S, left) and duration (D, middle). Dotted line is the fitted power-law exponent. Log likelihood ratio tests for S (left, outset) and D (middle, outset). Data points are coloured by fish. Avalanche schematic demonstrating the calculation of avalanche size (s, left top) and duration (d, middle top) for single avalanche events. Coloured neurons represent active neurons at *t_x_*. (right) Estimation of exponent relation. Empirical β is estimated by plotting mean S against D and fitting a linear regression line (dotted line). DCC is calculated as the absolute difference between empirical and predicted β. (B) DCCs plotted for each fish against each null model (dotted line = critical threshold of DCC < 0.2). (C) Branching ratio σ plotted for each fish against each null model (dotted line = critical value of σ~1). (top) Avalanche schematic demonstrating calculation of σ. (D) The quantity *r*(*d*) follows an approximate power-law with exponent *η* (dotted-line). (top) Schematic demonstrating estimation of correlation (*r*) as a function of distance (*d*). Magenta neuron is the neuron of interest. Other neurons are coloured by their correlation *r* to magenta neuron. * = p< 0.01

However, power-laws can emerge in non-critical systems. A more robust marker of criticality is exponent relation, which can distinguish between critical and non-critical regimes that produce power-laws (Ma et al., 2019; Touboul & Destexhe, 2017). Exponent relation dictates that at criticality the power-law exponents for avalanche size and duration can be predicted by a third exponent β (See Figure 2A). We use the deviation from criticality coefficient (DCC, see Methods) to assess exponent relation (Ma et al., 2019). Our datasets exhibit close to predicted β, suggesting the presence of near-critical dynamics (DCC = 0.13 ± 0.04) (See Figure 2B). In order to understand whether such relationships could have been generated by a random system, we created 3 null models: *spatial, correlation* and *autocorrelation* nulls.

With these models we were able to assess whether randomly generated neuronal activity, resulting from shuffling spatial structure, cell-cell correlation and autocorrelation respectively, would also generate critical DCCs. Importantly, exponent relation was significantly less well preserved in *spatial* nulls (DCC = 0.31 ± 0.02, t = −4.80, p < 0.001) compared to empirical data, but not *correlation* (DCC = 0.15 ± 0.04, t = −0.48, p = 0.64) and *autocorrelation* (DCC = 0.21 ± 0.05, t = −1.60, p = 0.15) nulls (see Figure 2B). This suggests that empirically observed exponent relations emerge due to the spatial structure of neural dynamics, rather than from random activity.

Another hallmark of criticality is a branching ratio (σ) close to 1. A branching process describes how activity is passed from an ancestor to its descendants, and only with critical branching (σ~1) can avalanches span all scales (Harris, 1963). We find σ close to 1 in our data (σ = 0.93 ± 0.03) further suggesting near-critical dynamics in spontaneous activity (See Figure 2C). Values of σ slightly below 1 suggest the presence of a slightly subcritical state (Priesemann et al., 2014), however σ is likely to be underestimated from multiple ancestors due to the convergence of activity onto shared descendants (see Methods). Importantly, we find that empirical σ is significantly closer to unity than *spatial* (σ = 0.77 ± 0.03, t = 21.8, p <0.0001), and *correlation* (σ = 0.92 ± 0.03, t = 4.45, p < 0.01), but not *autocorrelation* nulls (σ = 0.93 ± 0.03, t = −0.28, p = 0.79) (See Figure 2C). This suggests that σ~1 is a feature of the underlying spatial structure and correlation in neural activity.

Long-range correlations between neuronal signals are a hallmark of critical systems, as at criticality the correlation length is maximal. Long-range correlations give rise to power-law scaling of neuron-neuron distance and correlation in resting-state activity (Expert et al., 2011; Lombardi et al., 2020). We calculated the correlation function *r*(*d*) as the Pearson correlation between the activity of all neuron pairs as a function of distance (see Methods and Figure 2D). We find that *r*(*d*) qualitatively follows a power-law relationship which is well approximated by linear regression (See Figure 2D). This suggests the presence of both short and long-range correlations which can support avalanches that span the full spatial scale of the system.

We find that spontaneous dynamics exhibit (i) power-law relationships of neuronal avalanches, (ii) exponent relations close to critical values, (iii) branching ratios ~ 1 and (iv) power-law scaling of neuron correlation and distance. Taken together, these findings provide evidence that with physiological EI balance, the brain operates in a near-critical regime.

### EI Imbalance Causes Neural Dynamics to Deviate from Criticality

In a next step, we wanted to test whether EI balance acts to regulate criticality *in vivo*. Given that a critical point separates distinct phases, even small changes to a regulating parameter should give rise to abrupt changes in the system’s behaviour (Binney et al., 1992). Accordingly, perturbing EI balance with both partial and complete inhibitory blockade should cause dynamics to significantly deviate from criticality. Thus, we compared critical statistics following both *5mM* and *20mM PTZ* administration (See Figure S2).

We first compared avalanche exponents, to determine if perturbing EI balance alters cascading dynamics. Interestingly, exponents significantly decreased in both the *5mM* (Size: τ = 2.65 ± 0.10, w = 0, p < 0.01; Duration: α = 3.34 ± 0.12, w = 2.0, p < 0.01) and *20mM* conditions (Size: τ = 2.56 ± 0.05, t = 4.59, p < 0.01; Duration: α = 3.30 ± 0.11, t = 4.06, p < 0.01), compared with spontaneous activity (Size: τ = 2.98 ± 0.13; Duration: α = 3.58 ± 0.14) (See Figures 3B & S4). This indicates an increased frequency of larger and longer avalanches during seizures, suggesting a role of EI balance in shaping avalanche dynamics.

**F3.**
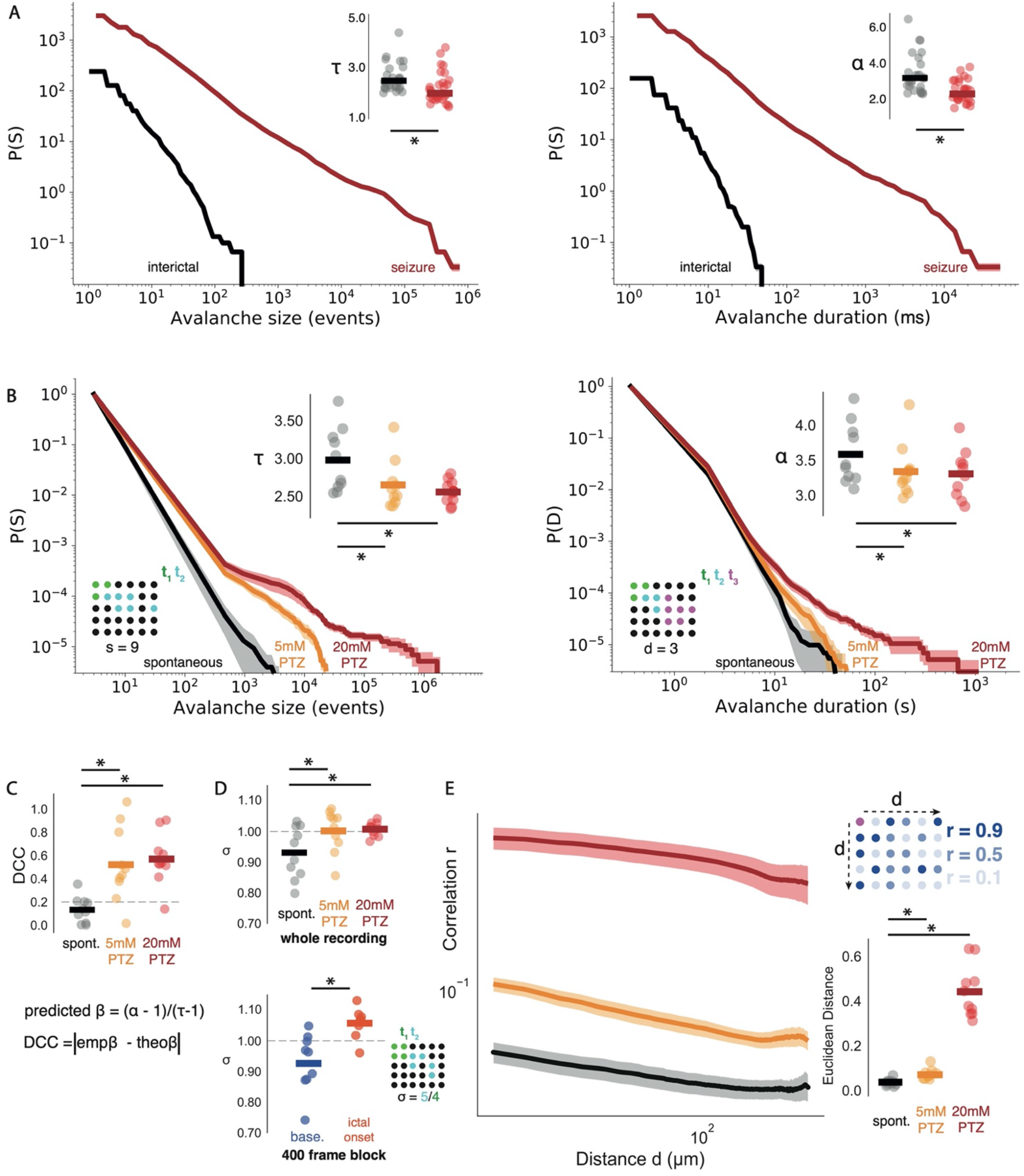
EI imbalance causes dynamics to deviate from criticality. (A)EEG data complementary cumulative distribution functions for avalanche size (S, left) and duration (D, right), comparing mean distributions across datasets. Shaded regions represent the standard error. Avalanche exponents compared for S (τ, left, top) and D (α, right, top) in interictal (black bar = mean), and *seizure* (red bar = mean) conditions. (B) Same as (A), for calcium imaging data with *spontaneous, 5mM PTZ* and *20mM PTZ* conditions. (C) DCC values plotted for each dataset. (D) σ plotted for each dataset across 30 minute (top) and 400 frame state transition blocks (bottom). (E) (left) The function *r*(*d*) compared across datasets. (right) Euclidean distance from fitted power law plotted for each dataset. * = p< 0.01

We were also interested in understanding whether EI imbalance-induced changes to avalanche dynamics were conserved across brain scales and recording modalities. Importantly, we find that focal-onset seizures in epilepsy patients give rise to concordant decreases in avalanche exponents (Size: median τ = 1.94, 1.38-3.92, U = 3.71, p < 0.001; Duration: median α = 2.25, 1.47-3.77, U = 4.24, p < 0.001), when compared with inter-ictal periods (Size: median τ = 2.52, 1.99-5.9; Duration: median α = 3.16, 2.26-11.64) (See Methods & Figure 3A). Therefore, PTZ-induced seizures in zebrafish exhibit analogous changes to cascading dynamics as found in human epileptic seizures measured with EEG.

Next, we assessed the sensitivity of exponent relation to different PTZ doses. We found that perturbing EI balance causes a divergence of the observed exponent relation from its predicted value, with both *5mM PTZ* (DCC = 0.52 ± 0.10, t = −3.24, p = 0.01) and *20mM PTZ* (DCC = 0.57 ± 0.07, t = −5.70, p < 0.001) causing significant increases in DCCs (See Figure 3C). This suggests a loss of self-similarity in avalanche dynamics due to the emergence of large avalanches that grow quickly in time, meaning that a single exponent cannot capture the relationship between size and duration across scales. Therefore, altering EI balance causes a deviation from a near-critical state in whole brain dynamics.

Next, we assessed the sensitivity of the branching ratio σ to changes in EI balance. We find a significant increase in both the *5mM PTZ* (σ = 1.00 ± 0.02, t = −3.79, p <0.01) and *20mM PTZ* conditions (σ = 1.01 ± 0.01, t = −3.11, p < 0.017), suggesting a key role of EI balance in shaping σ (See Figure 3D). We note that our seizure recordings contain both periods of ictal activity and post-ictal depression, and therefore σ closer to unity could arise due to averaging effects rather than critical dynamics. To evaluate this possibility we also compared σ during shorter windows of *baseline* and *ictal-onset* periods (see Methods).

Using this approach, we find that σ shows a greater magnitude increase beyond 1 during seizure onset (*baseline*: σ = 0.94 ± 0.03, *ictal-onset*: σ = 1.05 ± 0.02, t = −3.41, p < 0.01) (See Figure 3D). Thus, perturbing EI balance causes σ to increase beyond values suggestive of near-critical dynamics at rest, suggesting that EI balance shapes critical dynamics.

Finally, we assessed the sensitivity of correlation function power-laws to changes in EI balance. Interestingly correlation functions were less well approximated by power-laws in both the *5mM* (Euclidean distance from power-law = 0.07 ± 0.01, w = 1.0, p <0.01) and *20mm PTZ* conditions (Euclidean distance = 0.44 ± 0.04, w= 0.0, p <0.01) compared with spontaneous datasets (Euclidean distance = 0.04 ± 0.00) (See Methods, Figures 3E & S5). Therefore, EI imbalance saturates the system with excessive correlation thus disrupting power-law relationships that are a hallmark of criticality.

We find that perturbing EI balance causes a loss of critical statistics with i) changes to avalanche exponents, ii) a departure from exponent relationships, iii) increases in σ, and iv) a loss of correlation function power-laws. That even small changes to EI balance resulted in dramatic shifts in critical statistics, suggests that spontaneous dynamics may be finely tuned by excitatory and inhibitory connections near to a transition separating distinct phases of activity.

### Reorganisation of Network Topology in Network Model Pushes the Brain Away from Criticality

To understand how changes in synaptic EI balance shape the collective behaviour of wholebrain networks, we modelled the brain as a recurrent network of excitatory leaky-integrate-and-fire neurons. We were particularly interested in the network mechanisms supporting the emergence of pathological state transitions. Accordingly, we fit our model to 3 avalanche datasets comprising *baseline, pre-ictal* and *ictal-onset* periods (See Methods). We investigated 3 parameters that could be altered by synaptic EI imbalance (See Figure 4): 1) network topology (*m*) - the density of effective connections between neurons, 2) network geometry (*r*) - the distribution of edge weights between neurons, and 3) intrinsic excitability (*v_th_*) - the excitability of individual neurons. Thus in this model, we are subsuming effects of changes in local inhibitory connectivity into changes in the effective connectivity patterns between excitatory neurons, to provide a conceptual link between micro-scale biology and network-based analyses typically applied in clinical neuroscience (Bullmore & Sporns, 2009). Using all 3 parameters, we generated good model fits to *baseline* (*m* = 7, *r* = 5, *v_th_* = 20, cost = 0.113), *pre-ictal* (*m* = 6, *r* = 0, *v_th_* = 16, cost = 0.176), and *ictal-onset* data (*m* = 31, *r* = 1, *v_th_* = 17, cost = 0.120) (See Figure 4A). To compare the importance of each parameter we explored models containing a subset of parameters free for fitting, while keeping others fixed.

**F4.**
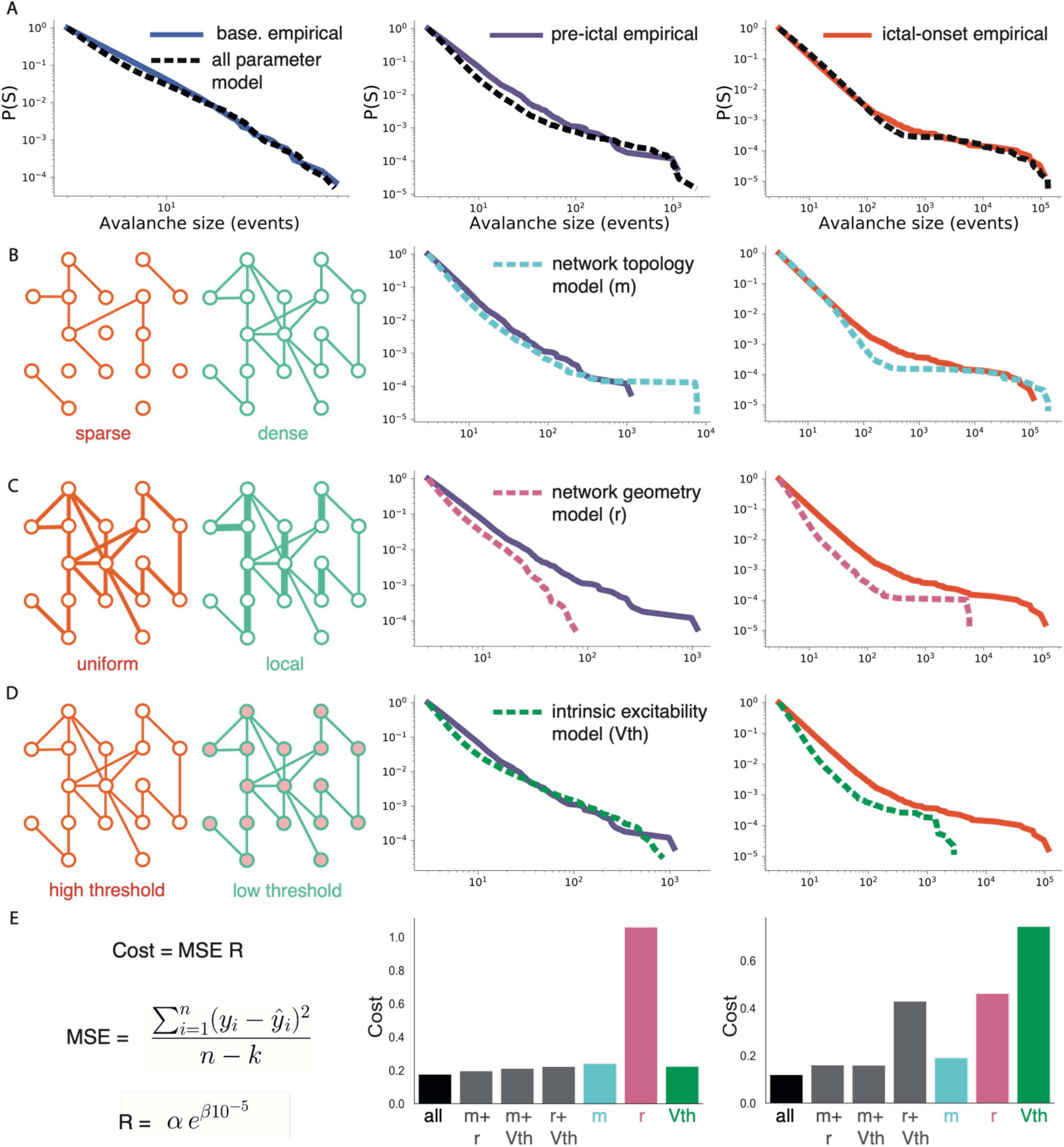
Reorganisation of network topology drives generalised seizures. (A) Avalanche distributions from *baseline, pre-ictal* and *ictal-onset* periods, alongside best model fits using all parameters. (B) (left) Network topology schematic for sparse and dense networks. Best model fits using *m* only to *pre-ictal* (middle) and *ictal-onset* data (right). (C) (left) Network geometry schematic for uniform and local weight scaling (edge thickness = synaptic strength). Best model fits using *r* only to *pre-ictal* (middle) and *ictal-onset* data (right). (D) (left) Intrinsic excitability schematic for high and low threshold to spike networks (red node = closer to spike). Best model fits using *v_th_* only to *pre-ictal* (middle) and *ictal-onset* data (right). (E) (right) Model comparison for 3 parameter models (black), 2 parameter models (grey) and single parameter models, for *pre-ictal* (middle) and *ictal-onset* data. (left) The cost function used for model comparison was a regularised mean squared error term (See Methods).

To model the emergence of the *pre-ictal* state, we fixed parameters to the best *baseline* model fit while allowing subsets of parameters to vary. This approach demonstrated that *v_th_* provided the best fit to *pre-ictal* data (*v_th_* = 19, cost = 0.223) (See Figure 4D). Altering *m* alone showed avalanche distributions with excessive heavy tails (*m* = 15, cost = 0.241), while *r* alone was unable to generate sufficiently heavy tails (*r* = 6, cost = 1.06) (See Figures 4B & 4C). However, we find that the model with all 3 free parameters provides the best *pre-ictal* fit, while models with any combination of 2 free parameters provide better fits than *v_th_* alone (See Figure 4E). Therefore the *pre-ictal* state likely emerges from small, non-specific changes to network topology, network geometry and intrinsic excitability.

To model the emergence of the *ictal-onset* state, we fixed parameters to the best *pre-ictal* model fit while allowing subsets of parameters to vary. Strikingly, only *m* provided a reasonable fit to *ictal-onset* data (*m* = 30, cost = 0.190) (See Figure 4B). Both *r* and *v_th_* alone failed to generate sufficiently heavy tails (*r* = 7, cost = 0.465; *v_th_* = 15, cost = 0.744) (See Figures 4C & D). In fact, while model fits with all 3 parameters provided the best overall fit, *m* alone provided a better fit than the model with both *r* and *v_th_* free to vary (See Figure 4E). This suggests that a reorganisation of network topology through an increase in edge density, best explains the loss of criticality that emerges with the onset of generalised seizures.

### Seizure Dynamics are Chaotic which Impairs Network Response Properties

The EI-associated changes in network topology characterised above will impact dynamics and ultimately information processing (Ju et al., 2020). In particular, σ > 1 suggests the emergence of a supercritical state (Meisel, 2020) which should impair network response properties (Shew et al., 2009). For such a state to arise we would also expect the emergence of chaos, as supercritical transitions *in silico* give rise to chaotic dynamics (Haldeman & Beggs, 2005). Thus to confirm the presence of supercritical dynamics in seizures, we first estimated the chaoticity of generalised seizure dynamics.

We approximated the largest Lyapunov exponent (*λ*) which estimates the divergence of nearby trajectories in phase space, a defining feature of chaos (See Methods & Figure 5A). As predicted, generalised seizures are significantly more chaotic (*20mM PTZ*: λ = 0.0034 ± 0.0001, t = −10.7, p <0.0001), than *spontaneous* dynamics (*λ* = 0.0024 ± 0.0001) (See Figure 5C). A more positive *λ* suggests that closeby trajectories will grow further apart over time, demonstrating a heightened sensitivity to initial conditions in the seizure state.

**F5.**
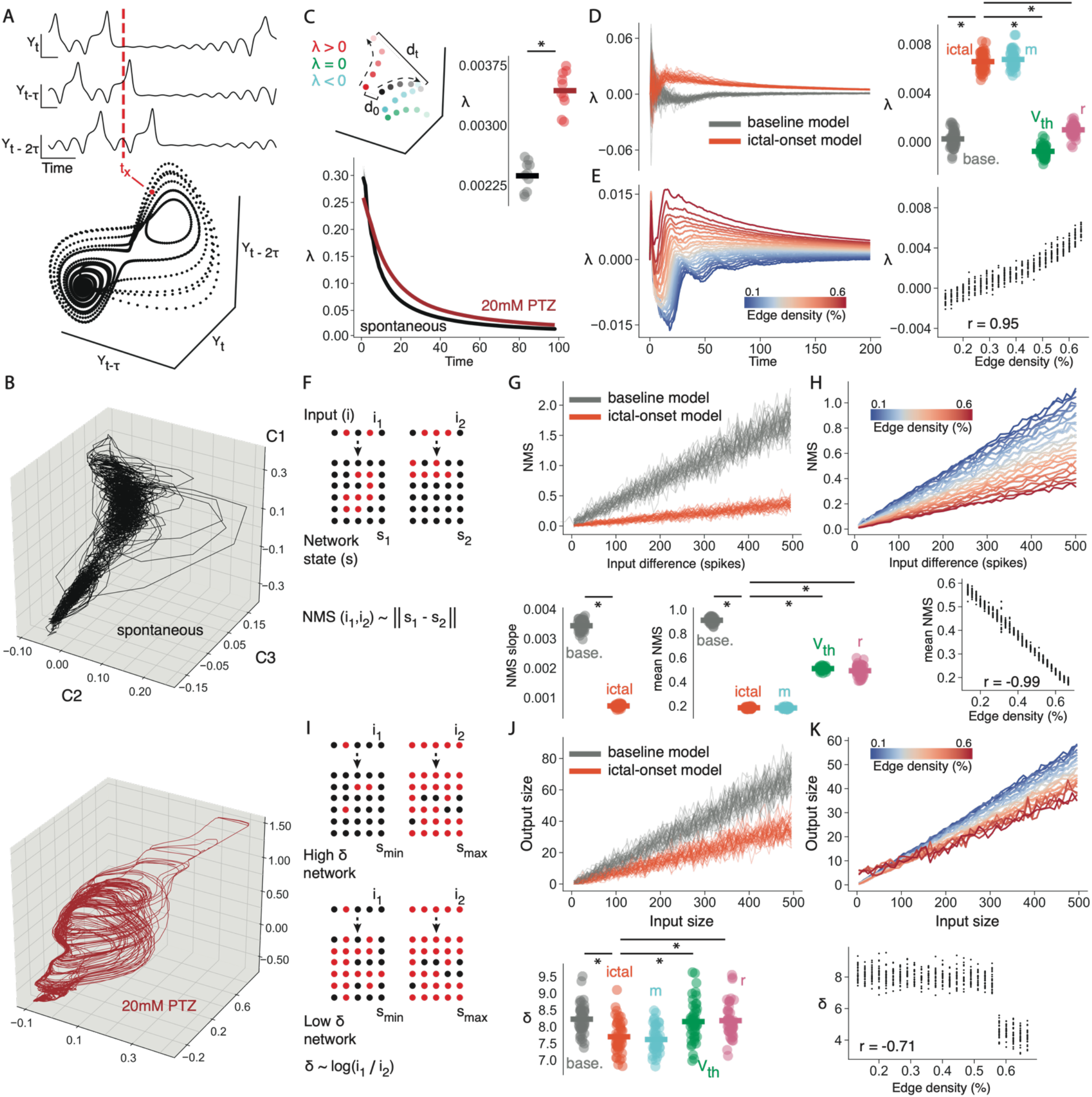
EI-imbalanced dynamics are supercritical. (A) (top) A single variable (*Y*) can be used to reconstruct an attractor that is topologically equivalent to the full system, by using a series of delayed variables (*Y, Y_t-τ_, Y_t-2τ_…, Y_t-(m-1)τ_*) of delay *τ* and dimension *m*. (bottom) Embedding each lagged variable into state space provides the reconstructed attractor, where *tx* is the position in state space at time *x* corresponding to the dotted red line in the above panel. (B) Isomap 3d embeddings of reconstructed attractors using lagged coordinate embedding for a representative fish. (C) (top) Schematic outlining the meaning of different *λ* values. Each colour represents the trajectory over time for a specific initial point along the attractor (high to low brightness represents movement in time). *λ* is the ratio of the difference between 2 points at the start (d0) and at *t* (dt). *λ* for each trajectory is calculated against the black trajectory. *λ* > 1: chaotic (red), *λ* < 1: stable (blue), *λ* = 1: neutral (green). (bottom) *λ* for *spontaneous* and *20mM PTZ* conditions. (right) Mean *λ* compared across *spontaneous* (black bar = mean) and *20mM PTZ* conditions (red bar = mean). (D) (left) *λ* compared for *baseline* and *ictal-onset* full parameter models. (right) *λ* compared for *baseline* and *ictal-onset* full parameter models and single parameter models. (E) (left) *λ* for increasing *m*, ranging from *pre-ictal* values to *ictal-onset* values. Each line represents the mean *λ* over time for each model over 50 simulations. (right) Correlation between *m* and mean *λ*. (F) For 2 similar inputs (*i_1_* and *i_2_*) onto the network, NMS is the Euclidean distance between the two corresponding network states (*s_1_* and *s_2_*). (G) (top) NMS as a function of input difference compared for *baseline* and *ictal-onset* full parameter models. (bottom) NMS slope and mean NMS compared for *baseline* and *ictal-onset* full models and single parameter models. (H) (top) NMS as a function of input difference for increasing *m*. (bottom) Correlation between *m* and mean NMS. (I) For different inputs, *δ* is the log ratio of the input sizes (*i_1_* and *i_2_*) that give rise to the largest and smallest network responses (*s_max_* and *s_min_*). (J) (top) Output size as a function of input size compared for *baseline* and *ictal-onset* full parameter models. (bottom) *δ* compared for *baseline* and *ictal-onset* full models and single parameter models. (K) (top) Output size as a function of input size for increasing *m*. (bottom) Correlation between *m* and *δ*. * = p< 0.01.

In line with our empirical data we also find that *ictal-onset* models are significantly more chaotic than *baseline* models (*baseline:* λ = 0.0002 ± 0.0001, *20mM PTZ:* λ = 0.0065 ± 0.0001, t = −45.5, p < 0.0001), using models with all parameters free to vary (See Figure 5D). Therefore during generalised seizures the brain enters a chaotic state, as expected for a supercritical network. The fact that spontaneous dynamics are neutral (*λ*~0) as expected near a bifurcation point, provides further evidence for near-critical dynamics at rest.

We also demonstrate that a reorganisation to network topology drives this chaotic transition, as only the *m* model produces *λ* matching the full *ictal-onset* model (0.0067 ± 0.0001, U = 1108.0, p = 0.16), while *r* (0.0010 ± 0.0001, U = 0.0, p < 0.0001) and *v_th_* (−0.0010 ± 0.0001, U = 0.0, p < 0.0001) models fail to produce chaotic dynamics (See Figure 5D). Furthermore, we find a significant positive correlation between *m* and *λ* (r = 0.95, p < 0.0001) (See Figure 5E). Therefore, increasing edge density drives the transition from a critical to a supercritical state during seizures.

To confirm that supercritical dynamics impair brain function, we assessed the impact of generalised seizures on 2 network response properties that are maximised at criticality: network-mediated separation (NMS), and dynamic range. NMS is the property of a network to encode distinct inputs with distinct network states, enabling similar inputs to be separated and discriminated by a readout function (Schrauwen et al., 2008) (See Methods & Figure 5F). For a network to reliably represent input differences of varying magnitude, NMS should increase proportionally to the difference between input signals. As predicted, *baseline* networks assume higher NMS values across the range of input sizes compared with *ictal-onset* networks (*baseline*: mean NMS = 0.91 ± 0.01, *ictal-onset:* 0.18 ± 0.00, U = 556991.0, p < 0.0001). Furthermore, *ictal-onset* networks exhibit a significantly more shallow NMS curve (*baseline:* slope = 0.0034 ± 0.0000, *ictal-onset:* slope = 0.0007 ± 0.0000, t = 109.5, p < 0.0001), in line with edge-of-chaos computing predictions for chaotic networks (See Figure 5G) (Maass et al., 2002). Therefore, seizures cause both a reduction in NMS and a reduced sensitivity of NMS to input size differences, indicating more similar network responses to large and small inputs. As above, we find that only the *m* model produced NMS values that matched the full model (*m*: 0.18 ± 0.00, U = 3081443.4, p = 0.20; *r*: 0.49 ± 0.01, U = 1104945.0, p < 0.0001; *v_th_*: 0.51 ± 0.01, U = 1546742.0, p < 0.0001) (See Figure 5G). We also find a significant negative correlation between *m* and NMS values (mean NMS: r =- 0.99, p < 0.0001, NMS slope: r = −0.99, p < 0.0001), indicating a link between increased edge density and reduced NMS (See Figure 5H).

The dynamic range (*δ*) is a measure of the range of input sizes that a system can represent (See Methods & Figure 5I). We find that *ictal-onset* networks exhibit significantly lower *δ* than *baseline* networks (*baseline:* 8.22 ± 0.06, *ictal-onset:* 7.70 ± 0.07, t = 5.83, p < 0.0001). This suggests that seizure networks exhibit a reduced capacity to represent a wide range of inputs, as expected in a supercritical state. Once again, only the *m* model matched the full *ictal-onset* model for *δ* (*m*: 7.62 ± 0.05, t = −0.95, p = 0.34; *r*: 8.18 ± 0.07, U = 593.0, p < 0.0001; *v_th_*: 8.15 ± 0.08, t = 4.28, p < 0.0001) (See Figure 5J). The parameter *m* was also significantly negatively correlated with *δ* (r = −0.71, p < 0.0001), although we note a non-linear relationship (See Figure 5K).

Therefore, our model demonstrates that chaotic dynamics impair the optimal response properties of critical networks, resulting in reductions in NMS and dynamic range. These changes are driven by a reorganisation to network topology during generalised seizures. We conclude that seizure dynamics are consistent with a chaotic, supercritical system away from criticality.

### Seizures Cause a Loss of Metastability Due to the Emergence of Sticky States

Supercritical networks should give rise to both impaired network response properties and pathological spontaneous dynamics. A defining feature of spontaneous activity at criticality is metastable dynamics (Haldeman & Beggs, 2005; Wildie & Shanahan, 2012), a regime in which brain activity endlessly transitions between a series of unstable fixed points, or metastable states (Cocchi et al., 2017). At criticality the number of metastable states is maximal, which can promote flexible transitions (Cabral et al., 2011). Therefore, to understand whether seizures hinder metastable dynamics, as predicted for a supercritical state, we compared the total number of metastable states in our data (See Methods). Interestingly, we find that generalised seizures exhibit significantly fewer metastable states than spontaneous dynamics (*spontaneous:* 20.9 ± 3.30, *PTZ 20mM:* 11.9 ± 2.31, w = 1.0, p < 0.01) (See Figure 6B). This indicates that seizures limit the diversity of states the brain can enter into, as predicted with supercritical dynamics (Haldeman & Beggs, 2005).

**F6.**
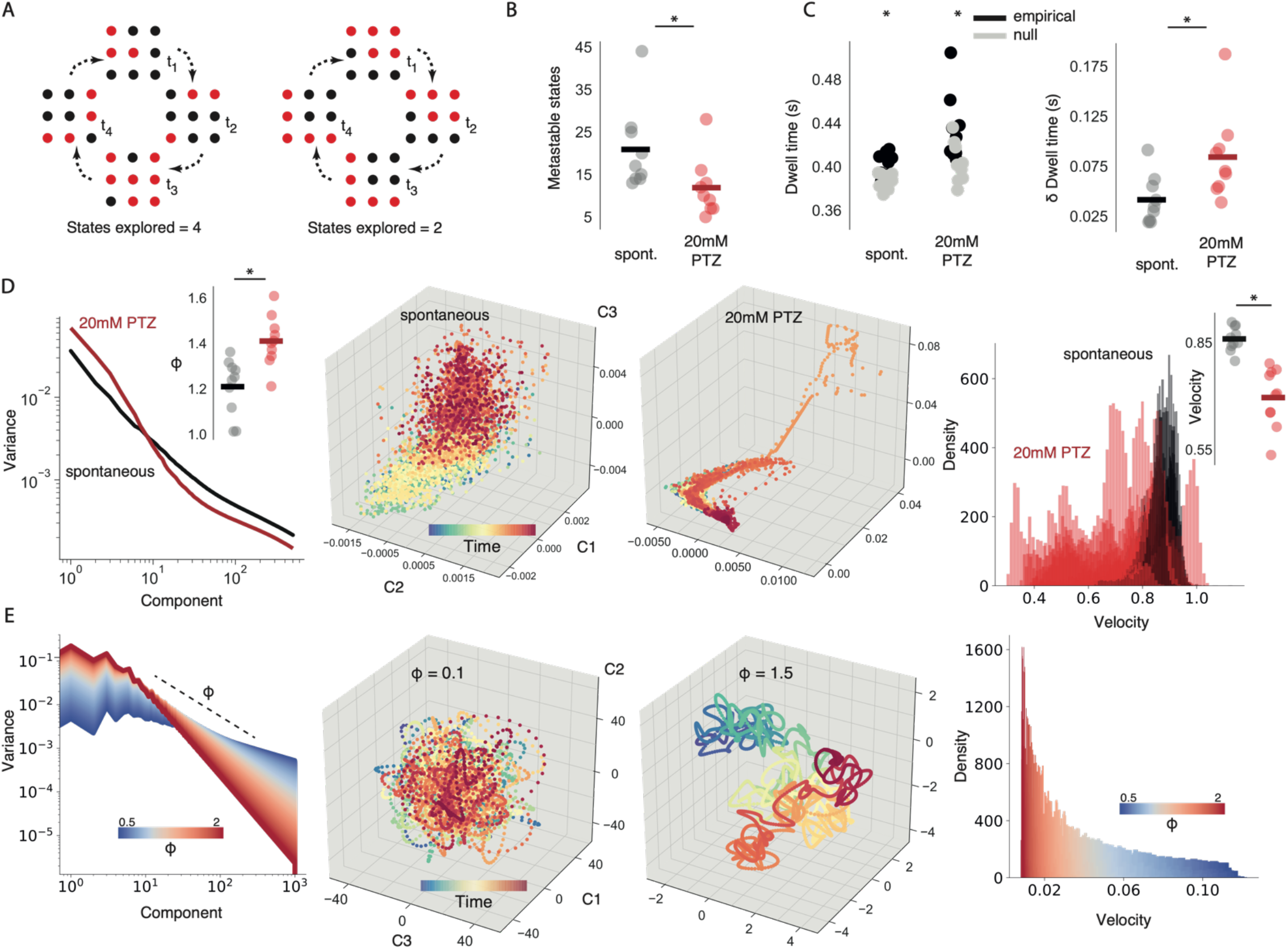
EI imbalance causes a loss of metastability. (A) A critical system can explore a greater subset of its possible brain states (left), while a non-critical system will explore a more limited subset (right). (B) Number of metastable states compared across *spontaneous* (black bar = mean) and *20mM PTZ* conditions (red bar = mean). (C) (left) Mean dwell time compared across conditions, plotting both *empirical* and *null* model datapoints. (right) Change in dwell time from null models compared across datasets. (D, left) Mean eigenspectrum function plotted across conditions. (outset) Eigenspectrum slope *ϕ* plotted for each dataset. (D, middle) 3d Isomap embedding of reconstructed attractor for an example fish. (D, right) State space velocity probability densities plotted for all fish, comparing *spontaneous* and *20mM PTZ* conditions. (outset) Comparison of mean velocity across datasets. (E, left) Simulated eigenspectrum function plotted for increasing *ϕ*. (E, middle) Random projection of eigenspectra into state space for different *ϕ*. (E, right) State space velocity probability densities plotted as a function of *ϕ*. * = p< 0.01

To understand how seizures might reduce metastability we hypothesised that high population covariance, driving the increase in edge density that we observe *in silico*, could constrain dynamics to fall into a limited subset of states. To test this we use the eigenspectrum function which defines the amount of variance *v*(*n*) explained by the *n^th^* principal component, thus providing a measure of multidimensional correlation. The eigenspectrum slope significantly increases during generalised seizures (*spontaneous: ϕ* = 1.20 ± 0.04; *20mM PTZ: ϕ* = 1.41 ± 0.03, t = −6.05, p < 0.001). This indicates greater variance captured in the first few components and increased population covariance during seizures (See Figure 6D). Interestingly, we note a direct relationship between *ϕ* and the velocity of the underlying dynamics (See Methods). As *ϕ* increases, due to greater variance captured in earlier components, population dynamics exhibit slower transitions in state space. To visualise this, we simulate eigenspectra power-laws and randomly project them into 3-dimensional space using a previously developed method (Stringer et al., 2019, See Methods & Figure 6E). In line with this, we find significantly slower dynamics in state space during generalised seizures compared with spontaneous activity (*spontaneous:* velocity = 0.86 ± 0.01, *20mM PTZ: velocity* = 0.70 ± 0.03, w = 0.0, p < 0.01) (See Figure 6D). Slower dynamics occur due to earlier components dominating the variance, such that the variance of reconstructed trajectories in state space is driven by only a few key modes. Slower transitions in state space suggest the emergence of sticky states which are difficult to leave. This could explain the loss of metastability, as sticky dynamics prevent the flexible transitioning into and out of the full dynamic repertoire.

For such sticky states to arise, we would expect longer times spent in each metastable state. In line with this, we find significantly increased dwell times during generalised seizures (*spontaneous:* dwell time (s) = 0.40 ± 0.01, *20mM PTZ*: dwell time (s) = 0.43 ± 0.01, w = 1.0, p < 0.01) (See Figure 6C). However, longer dwell times could occur by chance in systems with fewer states. Therefore we calculated null models for each system, by evaluating the dwell time expected by chance. We find that the dynamics for spontaneous and generalised seizure periods are non-random, with longer dwell times than expected by chance (*spontaneous:* w= 0.0, p < 0.01, *20mM PTZ*: w = 0.0, p < 0.01). Importantly, generalised seizures show significantly longer dwell times even when accounting for the fewer available states (*spontaneous*: δ dwell time (s) = 0.02 ± 0.01, *20mM PTZ*: δ dwell time (s) = 0.03 ± 0.01, w = 2.0, p < 0.05) (See Figure 6C).

Therefore, not only is metastability reduced away from criticality in line with a supercritical state, but alterations to population covariance give rise to slower dynamics and longer dwell times, suggesting the emergence of sticky dynamics. Such dynamics would likely prevent flexible responses to inputs and impair the optimal exploration of semi-stable states required for scale-invariant dynamics at criticality.

## Discussion

### Identifying mechanisms supporting criticality

Criticality describes a system in which dynamics are organised near to a phase transition, between order and disorder. An open question is how the brain can maintain dynamics at this balancing point, given the limited parameter range that defines a bifurcation region in a quasi-critical system (Bonachela et al., 2010). Maintaining critical dynamics in the brain would require adaptive feedback mechanisms between system state and regulating parameters (Chialvo et al., 2020; Sornette et al., 1995). This points towards homeostatic plasticity, which acts to regulate firing rates around a fixed point following changes to excitability (Tetzlaff et al., 2011; Zeraati et al., 2021). Key to the regulation of firing rates is the maintenance of appropriate excitation and inhibition (Turrigiano & Nelson, 2004), through activity-dependent changes to GABA (Kilman et al., 2002), NMDA and AMPA expression (Lissin et al., 1998). Given that homeostatic plasticity serves to regulate EI balance, the maintenance of EI balance could serve as a control mechanism for criticality.

Our data suggests that EI balance shapes critical dynamics *in vivo*. This is supported by *in vitro* studies which have shown that altering both inhibition and excitation alters critical dynamics (Beggs & Plenz, 2003; Bellay et al., 2015; Pasquale et al., 2008; Shew et al., 2009). We build on this work by investigating the link between EI balance and criticality using thousands of neurons across the whole brain *in vivo*, compared to hundreds of neurons *in vitro*. The fact that whole-brain critical statistics deviate dramatically following inhibitory blockade, demonstrates the sufficiency of EI balance for criticality in whole systems with sensory input and intact network connectivity. Furthermore, we use robust indicators of criticality not previously applied to EI balance, such as exponent relation and correlation functions, to provide further evidence for the importance of EI balance for critical dynamics. Taken together our data strongly suggest a role for EI balance in supporting near-critical dynamics.

Importantly, we note that critical statistics are correlative and can appear in non-critical systems. Thus critical statistics should be interpreted with caution (Priesemann et al., 2009). Furthermore, while the tuning of EI balance alters critical statistics as expected for a regulating parameter, further evidence is required to demonstrate an active role of homeostatic EI balance mechanisms in self-organising criticality. We also cannot eliminate the possibility of downstream mechanisms being influenced by changes to EI balance that may instead regulate criticality. For example, network topology is important for criticality and will be influenced by EI balance changes (Bornholdt & Rohlf, 2000). Furthermore, regulating gap junctions also gives rise to a deviation from criticality (Ponce-Alvarez et al., 2018). Therefore, while EI balance clearly shapes criticality, it is likely that multiple network mechanisms converge to maintain the brain at a phase transition.

### Understanding epileptic seizures through the lens of critical systems

We demonstrate that EI-imbalance induced epileptic seizures exhibit dynamics expected for supercritical networks. In particular, our data exhibit increased σ > 1 during seizure onset and the emergence of chaos, as predicted at supercriticality (Haldeman & Beggs, 2005; Shew et al., 2009). However, the interpretation of exact values of σ from empirical data can give misleading representations of system state (Priesemann et al., 2014). Nonetheless, σ can be useful as a comparative measure across the same system under different conditions. These findings naturally corroborate the impaired response properties exhibited in our supercritical networks: highly connected and sensitive networks give rise to exponentially growing avalanches regardless of input size, limiting the ability of the network to discriminate inputs. Therefore impaired network-response properties could give rise to the reduced responsiveness seen during seizures (Inoue & Mihara, 1998). Furthermore, our data indicate that supercritical networks cause increased population covariance which reduces the number of metastable states. Given the evidence for reduced brain state diversity during a loss of awareness induced by general anaesthesia (Schartner et al., 2015; Wenzel et al., 2019), we hypothesise that impaired metastability may explain the loss of awareness that occurs during generalised seizures (Blumenfeld & Taylor, 2003). In this way, supercritical networks may give rise to brain dysfunction in epilepsy.

However, criticality is one of many regimes that could optimise brain function, and in fact both non-critical and even chaotic networks can optimise computation (Farrell et al., 2019; Touboul & Destexhe, 2017). Furthermore, we note that PTZ-induced seizures in larval zebrafish may not faithfully recapitulate the diversity of seizure dynamics in epilepsy. Nonetheless, the larval zebrafish has emerged as a useful model for the study of genetic and pharmacologically-induced epileptic seizures (Burrows et al., 2020; Diaz Verdugo et al., 2019; Liao et al., 2019; J. Liu et al., 2021).

In conclusion, whole-brain *in vivo* dynamics deviate from criticality due to EI imbalance, indicating a regulating role for EI balance in shaping criticality. Epileptic seizures share statistics analogous to a supercritical state, showing the dynamics and response properties of a system driven away from criticality into disorder. Therefore seizures impair brain function by deviating from the critical point into chaos.

## Acknowledgements

DRWB is supported by an MRC-Sackler PhD Fellowship. DSB received funding from the John D. and Catherine T. MacArthur Foundation. MPM received funding from a Wellcome Investigator Award (204788/Z/16/Z). RER received funding from the Wellcome Trust (209164/Z/17/Z). We thank John Beggs and Fabrizio Lombardi for their helpful comments for the manuscript.

## Author Contributions

DRWB conducted zebrafish experiments. BP and MF conducted human experiments. DRWB, GD & RER analysed the data. DRWB built the models. DRWB, MPM & RER designed the research. DRWB, MPR, DSB, MPM & RER wrote the manuscript.

## STAR Methods

### Resource Availability

#### Key Resources Table

| **Regent or Resource** | **Source** | **Identifier** |
| --- | --- | --- |
| Tg(elavl3:H2B-GCaMP6s) | Freeman et al., 2014 | ZDB-TGCONSTRUCT-141023-1 |
| Scanimage Software - Vidrio Technologies (image acquisition) | This paper | https://docs.scanimage.org/ |
| Suite2p (image registration and segmentation) | Pachitariu et al., 2017 | https://github.com/MouseLand/suite2p |
| hidden markov model (calcium transient estimation) | Diana et al., 2019 | https://github.com/giovannidiana |
| Brian2 (Python network modelling software) | Stimberg et al., 2019 | https://github.com/brian-team/brian2 |

#### Lead Contact

Further information and requests for resources should be directed to the Lead Contact, Richard Rosch.

#### Materials Availability

Materials used to generate the study can be accessed on request from our lead contact.

#### Data and Code Availability

*Custom written python code can be accessed at: https://github.com/dmnburrows/criticality https://github.com/dmnburrows/avalanche_model https://github.com/dmnburrows/seizure_dynamics*

### Experimental Models and Subject Details

#### Larval zebrafish

Zebrafish larvae were raised at 28°C in Danieau solution on a day and night cycle of 12:12 hours. Transgenic zebrafish larvae, *Tg*(*elavl3:H2B-GCaMP6s*), expressing a nuclear localised genetically-encoded calcium sensor pan-neuronally were used for functional imaging experiments (gift from Misha Ahrens, Janelia Research Campus) (Chen et al., 2013; Freeman et al., 2014). To maximise optical transparency, *Tg*(*elavl3:H2B-GCaMP6s*) larvae were crossed with melanophore deficient (-/-) roy;nacre mitfa mutants (Lister et al., 1999). All imaging experiments were performed at 6 days post fertilisation (dpf). This work was approved by the local Animal Care and Use Committee (Kings College London), and was carried out in accordance with the Animals (Experimental Procedures) Act, 1986, under license from the United Kingdom Home Office.

#### Human participants

Anonymised intracranial EEG recordings from stereotactically implanted EEG electrodes (SEEG) were selected from the clinical database at Great Ormond Street Hospital, a national epilepsy surgery centre. Patients underwent SEEG recording with the aim to record seizures for presurgical evaluation of a pharmacoresistant focal epilepsy of presumed focal onset onset with or without visible lesion on clinical MR imaging. Ethical approval for the use of anonymised data from the clinical database for quantitative analysis was granted by the UK Health Regulatory Authority (IRAS 229772) and the Joint Research Office at the UCL-Great Ormond Street Institute of Child Health (Project ID 17NP05).

### Method Details

#### 2-photon calcium imaging

For imaging experiments, non-anaesthetised larvae at 6 dpf were immobilised in 2% low-melting point agarose (Sigma-Aldrich) and mounted dorsal side up on a raised glass platform that was placed in a custom-made Danieau-filled chamber. All imaging was performed on a custom built 2-photon microscope (Independent NeuroScience Services, INSS), which utilises a Mai Tai HP ultrafast Ti:Sapphire laser (Spectraphysics) tuned to 940nm. Objective laser power was at 15mW for all experiments. Emitted light was collected by a 16x, 1NA water immersion objective (Nikon) and detected via a gallium arsenide phosphide detector (ThorLabs). Scanning was performed by a resonance scanner (x-axis) and galvo-mirror (y-axis), with a piezo lens holder (Physik Intrumente) adjusting the z-plane (See Figure S1A). Volumetric data was collected across 10 focal planes at 15μm intervals, resulting in a frame rate of 2.73 Hz per volume with 4,914 volumes collected per imaging block (See Figure S1B). Larvae were left for 30 minutes in the light to allow the agarose to set and the fish to habituate to its environment.

To induce seizures we used the GABA_A_ antagonist pentylenetetrazole (PTZ), which elicits clonus-like convulsions and epileptiform discharges, which are removed with conventional antiseizure medication (Baraban et al., 2005). To capture EI balanced/imbalanced dynamics we recorded 3 x 30 minute consecutive imaging blocks for each fish: 1) *spontaneous* activity representing EI balance, 2) *5mM PTZ* causing EI imbalance giving rise to focal seizures, and 3) *20mM PTZ* causing EI imbalance giving rise to generalised seizures (See Figure S2). PTZ was added into the imaging chamber immediately after each session via a 1ml dose of PTZ suspended in danieau, after which point imaging was restarted. EI balanced dynamics in the *spontaneous* condition consisted of low correlation, local ensemble activity and rare, global activity which recruits large parts of the brain (See Figure S2A). EI imbalanced dynamics consisted of localised, high correlation ensemble activity in the *5mM PTZ* condition (See Figure S2B) and long-lasting, high-amplitude, synchronous activity recruiting most of the brain in the *20mM PTZ* condition (See Figure S2C). Data was collected and analysed in 10 fish.

#### 2 photon data processing

The use of nuclear localised GCaMP enables the segmentation of individual neurons according to somatic fluorescence. First recordings were visually inspected for the presence of z-drift, with datasets removed accordingly. Recordings were then corrected for drift in x and y planes by using the rigid registration algorithm on suite2p (Pachitariu et al., 2017). Briefly, this algorithm relies on phase correlation, which uses spatial whitening to compute the cross-correlation peaks between frames via fast Fourier transforms (Alba et al., 2015). We then use the segmentation algorithm on suite2p, which identifies highly correlated pixels in a spatial neighbourhood and models the fluorescence as the sum of neuronal activity, contamination from the surrounding neuropil and Gaussian noise. While the segmentation process is effective at identifying most of the cells in the recording, some segmentations may be false positives due to noise. To remove these non-cells, a false-positive detection algorithm was designed which relies on the assumption that true cells would show fluorescence spikes with a slower decay time than shot noise events, due to the decay time of GCaMP6s. The false-positive detection algorithm works as follows: 1) split the trace for each cell into 9 frame windows, 2) find the minimum fluorescence value across each window, 3) find the single maximum value across all minimum window values for each cell. This process creates a distribution of max-of-min fluorescence window values (a measure of the fluorescence decay speed), for all segmented cells across the brain. A threshold was then chosen to remove non-cells for each fish. This process enabled the accurate segmentation of ~9,000 neurons per fish.

In order to estimate calcium events from raw fluorescence we applied a hidden Markov model (HMM), developed by our lab and published in a previous paper (Diana et al., 2019). Briefly, the method models a neuron as a Bernoulli process with a latent variable *s_t_* representing its hidden on/off states. The fluorescence trace is decomposed into the sum of calcium transients *c_t_* with fixed decay constant *λ* and onset defined by *s_t_*, baseline activity *b_t_* and Gaussian noise. The algorithm iteratively maximises the probability of the latent state at time *t*, given the state of *t*-1 and the empirical observation of fluorescence value *x_t_*. This process enables the robust estimation of the onset of exponentially decaying calcium transients. The model requires the selection of parameter *q*, defining the event probability. This parameter’s value was chosen by visual inspection of which *q* gave the most accurate representation of calcium traces in our data (*q* = 0.59) (See Figure 1B).

For our analysis, data was compared across entire 30 minute imaging blocks (n=10) and shorter 400 frame periods, to examine generalised seizure state transitions (n=9). Generalised seizure transitions were defined as the abrupt appearance of whole-brain, synchronous cascades (See Figure S3). In order to define the onset of a state transition (in *20mm PTZ* imaging blocks) we required a clear separation between local ictal and globally synchronous ictal events: any brain recordings where the maximum mean whole-brain fluorescence was at least 4x greater than the minimum mean fluorescence were included in state transition analyses. This approach removed 1 dataset in which ongoing, synchronous activity occurred throughout the recording, making it difficult to identify specific generalised state transitions (See Figure S6B). For the remaining 9 datasets, to identify the beginning of generalised state transitions we used a 30 frame sliding window over the mean whole-brain fluorescence trace, calculating the maximum change from the start of the window and the subsequent 29 frames. The window with the highest difference was defined as the start of the generalised state transition. In order to compare dynamics across state transitions, we compared activity from 400 frames just before the generalised seizure (*pre-ictal*), the first 400 frames of the generalised seizure (*ictal-onset*) and 400 frames from a randomly selected segment of spontaneous recording (*baseline*) (See Figure S6A).

#### Human intracranial EEG recordings

For the analysis of human epileptic seizures, 30 patients were selected based on the availability of at least 3 seizures during the SEEG recording; as well as artefact-free baseline segments of corresponding duration that were >60 minutes away from ictal recordings. Seizure onsets were marked by a consultant clinical neurophysiologist specialising in presurgical evaluation of patients with focal epilepsy (FM) as part of routine clinical care. For each patient, data of 3×30 second windows of baseline awake EEG activity, and 3 clinically identified seizures were selected for analysis. Clinical SEEG data were recorded at a sampling frequency of 1024Hz. Raw SEEG data were re-referenced to an average reference, and filtered to a 1-250Hz broad frequency spectrum using a finite impulse response (FIR) filter using the window method with a Hamming filter window in the MNE (MEG and EEG analysis and visualization) toolbox in Python (Gramfort et al., 2013).

### Quantification and Statistical Analysis

#### Calcium imaging avalanche detection

Neuronal avalanches are defined as cascades of neural activity that propagate in space and time. Given that the underlying synaptic connectivity is not known, we must infer the flow of activity. To do this we adapt methods used in previous studies to estimate avalanche size and duration (Ponce-Alvarez et al., 2018; Tagliazucchi et al., 2012).

First, we assume that neuron *u* can only activate other neurons in the population *P* that lie within a local neighbourhood *N_u,v_* where *v* represents the closest *k*% of cells to *u*. This introduces a spatial constraint to avalanche propagation such that activity can only flow between putatively connected neurons, preventing disparate cascades combining into one large avalanche (See Figure 1). *k* was selected (0.16%) so that the mean neighbourhood radius (15.5um ± 0.78) lay within the neighbourhood range used to calculate critical statistics in previous studies (<30um).

We adapt the avalanche calculation algorithm defined by Ponce-Alvarez *et al.*, for single cell data (See Figure 1D). Briefly, an avalanche begins when at least 3 cells within a neighbourhood *N_u,v_*are active at *t_x_*, that were not part of an avalanche in the previous timestep. Here we label the neurons currently active at *t_x_* which belong to the avalanche as the set *A_x_* = {*a,b,c…z*}. All active neurons of *P* at *t_x_* connected to any of *A_x_* via a neighbourhood are included into the set *A_x_*.

At each subsequent step avalanche propagation iterates as follows:

1. If at least one neuron that was part of avalanche *A_x_* is also active at *t_x+1_* then the avalanche continues in time, forming the set *A_x+1_* = {*a,b,c…z*}.
2. Any neurons from *P* active at *t_x+1_* that are connected to any of *A_x+1_* via a neighbourhood are included into the set *A_x+1_*.

Once step 1 is no longer satisfied the avalanche terminates. Avalanches whose active neighbourhoods converge are grouped into a single avalanche. Avalanche size was calculated as the total number of calcium events during the avalanche. Avalanche duration was the number of time steps for which the avalanche was active.

#### Power-law distribution analysis

A common hallmark of critical systems is the presence of scale-invariant power-law distributions (Bak et al., 1987). In order to statistically evaluate the presence of power-law distributions in our data, we used an importance sampling approach to compare the likelihood of our data being generated by a power-law compared to a log normal distribution.

For a given power-law exponent λ, the likelihood of λ is given by

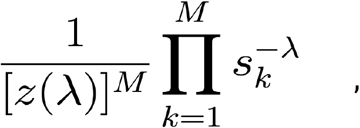

with avalanche size (or duration) defined from *s_1_, s_2_, s_3_…s_m_*.

The normalisation constant *z*(*λ*) is a function of the exponent

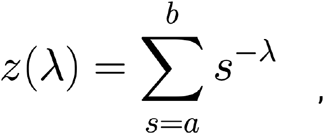

where *a* and *b* are the minimum and maximum avalanche cutoffs (*avalanche size: a* = 3, *b* = maximum observed for each fish, *duration: a* = 2, *b* = maximum observed for each fish).

The log likelihood (log L) is then defined as

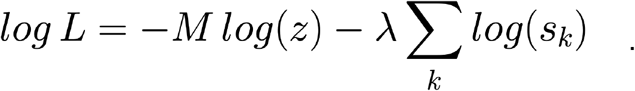

Log likelihoods were calculated across a range of sampled *λ*, which were weighted by the log probability of observing *λ_i_* from the prior and proposal distributions.

To be precise, the weight *w_i_* for exponent *λ_i_* is given by

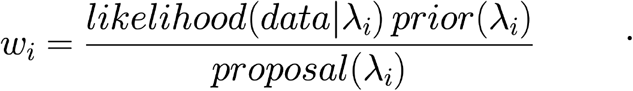

Marginal likelihoods (ML) were estimated as the empirical means of all the weights

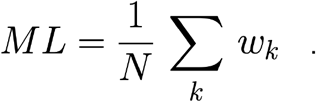

The likelihood ratio (LR) was calculated as

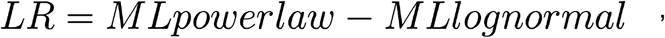

with positive values indicating the presence of power law distributions.

Avalanche exponents were calculated as the maximum likelihood of λ given the data. While some studies claim that critical systems should exhibit specific avalanche power-law exponents, such exponents vary according to temporal binning and avalanche-neighborhood radius (Beggs & Plenz, 2003; Ponce-Alvarez et al., 2018; Priesemann et al., 2013). In line with these prior reports, we find that avalanche exponents are dependent on an arbitrary choice of avalanche detection parameters, such as the probability of a calcium fluctuations being classified as a spike in the HMM (event probability), and % of closest neurons allocated to a neighbourhood in the avalanche detection algorithm (neighbourhood size) (See Figure S3A). Importantly, powerlaw fits to avalanche size and duration probability distributions are retained across all parameter combinations suggesting that power-law relationships are robust to parameter selection (See Figure S3B).

#### Exponent relation

The presence of power-laws alone is insufficient evidence of criticality, as they can arise from non-critical and random mechanisms (Miller, 1957; Stumpf & Porter, 2012; Touboul & Destexhe, 2017). A more reliable marker of criticality is exponent relation, which can separate out truly critical from chaotic, non-critical systems which generate power-law distributions (Touboul & Destexhe, 2017). Exponent relation states that for critical systems, avalanche size (*τ*) should scale as a power (*β*) of avalanche duration (*α*) (Tang & Bak, 1988). The exponent relation is defined as

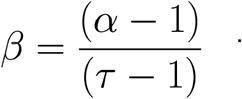

This relationship exists because critical systems are self similar, so a single exponent β can predict the relationship between τ and α across all scales of space and time. We use the previously developed deviation from criticality coefficient (DCC) to assess exponent relation (Ma et al., 2019). To calculate DCC we predict theoretical *β* using power law exponents estimated by maximum likelihood estimation for size (*τ*) and duration (*α*). We then estimate empirical *β* values from avalanche distributions by plotting the mean size <S> against duration and fitting exponents via linear regression. DCC is defined as the absolute difference between predicted and empirical *β* (See Figure 2A). We find that DCC values varied smoothly as a function of the event probability, with lower event probability giving rise to greater DCC values (See Figure S3D). Nonetheless, near-critical values (DCC < 0.25) were retained for event probabilities that accurately matched calcium transients (event probability = 0.57 - 0.61), regardless of neighbourhood size.

#### Branching Ratio

The branching parameter σ describes the propagation of activity over time, from ancestors to descendants. The quantity *σ* is defined as the average number of descendants from one ancestor (Harris, 1963). Critical systems should have a *σ* ~ 1, which enables avalanches that span all scales of space and time. We estimate *σ* from multiple ancestors similar to a previous study (Beggs & Plenz, 2003), as estimating *σ* from individual ancestors is not practical in our dataset given the large number of neurons active at any given timestep. The quantity *σ* was calculated as the mean ratio of descendants to ancestors at each time step over all avalanches as

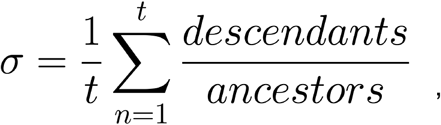

where *t* is the total number of avalanche propagation timesteps. Previous studies have demonstrated that σ is affected by temporal binning and subsampling (Priesemann et al., 2009). In line with this, we find that σ values vary according to both event probability and neighbourhood size (See Figure S3C).

#### Correlation Functions

Power-law relationships of neuron-neuron distance and correlation is expected at criticality. In order to test this hypothesis we calculated the correlation function *r*(*d*), where *r* indicates the Pearson’s correlation coefficient between two activity time series and *d* indicates the Euclidean distance between the two neurons producing the activity time series. First, neuron-neuron paired distances were placed in 200 bins, equally spaced across the range of neuron-neuron distances of all neurons in the brain. The quantity *r*(*d*) was then estimated as the mean correlation within each bin (See Figure 2D). Given that these functions are not probability distributions, we cannot perform the likelihood ratio tests described above. Thus to assess closeness of fit to power-laws we calculated the Euclidean distance between fitted power-law (fitted by linear regression) and the empirical data.

#### Null models

In order to test whether critical statistics emerge due to the spatiotemporal structure of neural activity rather than from random activity, we generated a series of null models. The first null model is the *spatial* null, which tests the hypothesis that critical dynamics emerge from spatially contiguous units. The *spatial* null is generated by randomly shuffling the locations of neurons in the brain, while retaining the temporal structure. The second null model is the *correlation* null, which tests the hypothesis that critical dynamics are generated by the correlation structure of neural activity. The correlation null was generated by circularly permuting the time series independently for each neuron, such that the correlation across neurons was lost, while retaining the autocorrelation structure. The third null model is the *autocorrelation null*, which tests the hypothesis that avalanche dynamics are generated by the integration of activity over time within neurons. The *autocorrelation* null was generated by randomly shuffling the binarised activity of all neurons in a dependent fashion, such that autocorrelation was lost but the cell-cell correlation retained. In order to compare critical statistics from empirical and null data, 50 nulls were randomly generated for each fish and for each null type, with avalanches calculated as above.

#### EEG avalanche analysis

The signal of both seizure and baseline recordings from each SEEG channel was z-scored against the baseline recording of that SEEG channel by subtracting the baseline mean and dividing it by the baseline standard deviation. Data were then binarised by identifying peaks that exceed a peak amplitude threshold parameter *p* (Arviv et al., 2016). Individual peaks across channels were grouped into a single avalanche if they occurred within a short time window Δ*t*. Thus, neuronal avalanches in the EEG data were defined as sequences of spatially distributed peaks of oscillatory activity. Statistical evaluation of the resultant distribution of avalanches was performed as described for zebrafish data above using a likelihood ratio (LR) of the data being derived from a power law vs. the data being generated by a log-normal distribution. A parameter sweep across values across frequency bands (broad 1-250Hz, delta 1-4Hz, theta 4-8Hz, alpha 8-15Hz, beta 15-30Hz, gamma 30-80Hz, and high gamma 80-250Hz), values for *p* (2-6 in 12 increments) and Δ*t* (2-20ms in 20 increments) was performed and for each resultant avalanche distribution in the baseline data, we evaluated the LR, demonstrating a likelihood ratio in favour of the baseline data being a power law distribution for the majority of parameter combinations (See Figure S3B). For subsequent analysis parameter values of *p* = 4 and Δ*t* = 5 were chosen and the analysis was performed on broadband (1-250Hz) filtered data. Avalanche statistics were compared across baseline (interictal) and seizure datasets. Power law exponents to the size and duration of individual avalanches were calculated as described for zebrafish data above, and their distribution was compared across baseline (interictal) and seizure data.

#### Model Fitting

To model avalanche dynamics in the brain we consider a network of leaky integrate-and-fire neurons with *N* excitatory neurons (*N* = 8990), and *E* external excitatory inputs (*E* = 1000) onto each neuron. The membrane voltage for neuron *i* is defined at subthreshold voltages by the differential equation

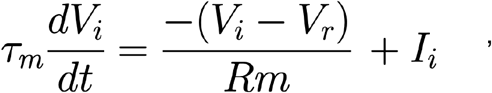

where *V_i_* is the membrane voltage, *τ_m_* is the membrane time constant (*τ_m_* = 20), *V_r_* is the resting membrane potential (*V_r_* = 0), *R_m_* is the resistance, and *I_i_* is the input current. For voltages beyond the voltage threshold *v_th_* the neuron fires a spike, at which point *V_i_* is reset to *V_r_*.

The input to neuron *i* at time *t* is defined by

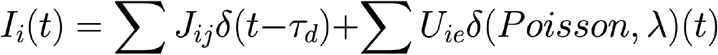

The first term describes the input from other neurons in the network, where *J_ij_* is the weighted adjacency matrix, between the presynaptic neuron *i* and postsynaptic neuron *j*. A spike from neuron *j* will affect neuron *i* after a synaptic delay *τ_d_* = 1, through Dirac’s delta function. The second term describes the input from the external current, where *U* is the weighted adjacency matrix between the presynaptic neuron *i* and the external neuron *e*. External input spikes are modelled as a Poisson point process with rate λ = 10. The quantities *U_ie_* for all connections were set to 0.1.

In order to construct a network with biologically realistic neuron-neuron distance distributions, we registered all datasets (10 fish, 3 conditions) to a standard space (Tabor et al., 2019). We then performed k-means clustering on the spatial locations of all neurons (*k* = 8990, the mean number of cells across all datasets), with resulting cluster centroids used as network node locations. Network connectivity was defined by the preferential-attachment algorithm (Barabási & Albert, 1999). The algorithm begins with *m* initial connected nodes and progresses by sequentially adding a new node that forms *m* connections with existing nodes, where the probability *p_i_* that each new node is connected to node *i* is proportional to the degree of node *i* defined as

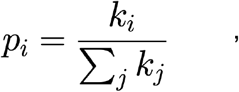

where *k_i_* is the degree of node *i* and *k_j_* is summed over all pre-existing nodes.

We defined 3 parameters of interest in our model (See Figure 4): 1) network topology - the density of effective connections between neurons defined by parameter *m*, 2) network geometry - the distribution of edge weights between nearby vs. distant neurons defined by parameter *r* and 3) intrinsic excitability - the excitability of individual neurons defined by parameter *v_th_*. Network topology was varied by altering *m* in the growth and preferential-attachment algorithm, thus changing the number of binary edges between all nodes. Network geometry defines the distributions of edge weights across the network. We wanted to probe the extent to which the network shows a preference for local versus uniform synaptic weights, i.e. stronger synaptic weights for shorter connections, or a uniform distribution across neuron-neuron distances. To vary network geometry smoothly we created a weight function *w*(*d*) which defines synaptic weight as a function of distance, where *w*(*d*) is defined

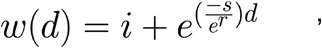

where *i* is the initial non-scaled synaptic weight (*i* = 1.2), *s* is the softening parameter that dictates the magnitude of exponential decay for the synaptic weight over distance (*s* = 0.1), *d* is the neuron-neuron distance and *r* defines the local versus uniform synaptic weight distribution (high *r* = weight scaling more uniform across distance, low *r* = weight scaling preference for closer neurons). Intrinsic excitability was represented as the spike threshold of each node in the network, defined by varying *v* for each neuron.

We simulated network dynamics for 4000 time steps. Spike data was then binned into 400 time bins, from which avalanche distributions were calculated as above. Model avalanche distributions were fit to empirical data generated from concatenated distributions for avalanche size of *spontaneous, pre-ictal* and *ictal-onset* datasets (n=9, 400 frames per dataset).

Avalanche simulations were run 9 times, and concatenated together to generate fits to empirical data.

To fit the model to our data we estimated the distance between empirical avalanche distributions and model avalanche distributions. The cost function was a regularised mean squared error term defined as

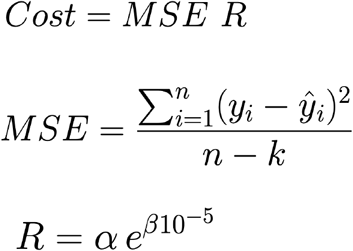

where *y_i_* = *P*(*S*) for each avalanche probability distribution bin, *n* is the number of avalanche probability distribution bins, *k* is the number of parameters, *β* is the difference in the number of non-empty bins between model and empirical data, and *α* scales the effect of the regularisation (*α* = 0.09).

We performed a grid-search across the range of parameter values which captured the full extent of empirical avalanche distribution shapes (*m* = 5 - 33), (*r* = 0 - 7), (*v_th_* = 15 - 20), exploring approximately 1,400 parameter combinations in total. This approach allowed us to find the best model fits using all 3 parameters for *spontaneous* (*m* = 7, *r* = 5, *v_th_* = 20, cost = 0.113), *pre-ictal* (*m* = 6, *r* = 0, *v_th_* = 16, cost = 0.176), and *ictal-onset* data (*m* = 31, *r* = 1, *v_th_* = 17, cost = 0.120) (See Figure 4A). In order to compare the importance of different model parameters we then explored a subset of parameters that were free to vary, while keeping others fixed. To explain the emergence of the *pre-ictal* state, we fixed parameters to the best *spontaneous* model fit while allowing subsets of parameters to vary freely for model fitting. To explain the emergence of the *ictal-onset* state, we fixed parameters to the best *pre-ictal* model fit.

#### Model response properties

In order to assess the ability of a network to separate out distinct inputs we measured the network-mediated separation (NMS) property. To calculate NMS we provided pairs of inputs, *a* and *b* separated by distance *z*. For each input, *x* nodes were randomly activated once the network had reached steady state at time *t* (4000 time steps). Network activity was binned into 10 time step windows, as above. In order to distinguish between the separation of network activity due to the network response to the input as opposed to the intrinsic variability in the network, we calculated a normalised NMS property as

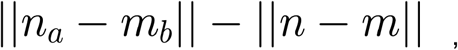

where ∥ ∥ denotes the Euclidean distance. The vectors *n_a_* and *n* are the binarised state vectors at time *t*, for identical instantiations of a network that received input *a* or received no input, respectively. The vectors *m_b_* and *m* are the binarised state vectors at time *t*, for identical instantiations of a network that received input *b* or received no input, respectively. To assess the slope of the NMS curve we performed linear regression on the mean NMS against input size difference.

In order to assess the range of information that the network can hold we measured the dynamic range (*δ*) of the network, adapting previous measures (Shew et al., 2009). To calculate *δ* we provided input to the network by randomly activating *x* neurons across a range of input sizes (*x* range = 5 - 500, stepsize = 10), once the network had reached steady state (4000 time steps).

Network activity was binned into 10 time step windows, as above. In order to separate out the network response to a given input from the ongoing network activity, the corresponding output size was calculated as

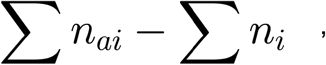

where *n_a_* and *n* are binarised state vectors for identical instantiations of a network that received input *a* or did not receive input at time *t*, respectively.

The quantity *δ* is then defined across the range of input sizes as

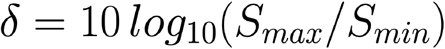

where *S_max_* and *S_min_* are the input sizes leading to 90th and 10th percentile over the range of output sizes, respectively.

#### Lyapunov exponent estimation

In order to test for the presence of chaotic dynamics in our empirical data, we use delay embedding theorem to reconstruct the attractor of the underlying dynamical system (Takens, 1981). The delay embedding theorem states that a reconstructed manifold constructed from a variable *Y_0_*, given by {*Y_0_*(*t*), *Y_0_*(*t - τ*),…, *Y_0_*(*t* - (*m - 1*)*τ*)} is topologically equivalent to the full dynamical system {*Y_0_*(*t*), *Y_1_*(*t*),…, *Y_m-1_*(*t*)}, where *m* is the dimension of the system and *τ* is the time lag (Takens, 1981) (See Figure 5A). We reconstructed the attractor using the first principal component of each dataset, as it captures the most variance in the system.

First, *τ* was calculated for each dataset as the *τ* which maximised the mutual information between {*Y_0_*(*0*), *Y_0_*(*1*), …, *Y_0_*(*n-1*)} and {*Y_0_*(*0 + τ*), *Y_0_*(*1+τ*), …, *Y_0_*((*n-1*) + *τ*)}. Specifically, for a given *τ* the mutual information was quantified by binning the time series and calculating

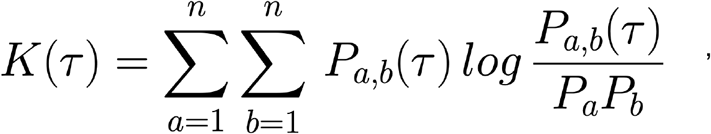

where *n* is the number of bins, and where *P_a_* and *P_b_* are the probabilities that an element lies in bin *a* and *b*, respectively. The quantity *P_a,b_*(*τ*) represents the probability that *Y_0_*(*x*) lies in bin *a* while *Y_0_*(*x+τ*) lies in bin *b*. We selected the *τ* which maximised the mutual information for each time series.

The embedding dimension *m* was estimated using a false nearest neighbours approach which assumes that at the correct dimension, nearest neighbours should be retained at higher dimensions. Points in state space were nearest neighbours for a given *m* if their distance was smaller than the standard deviation. False nearest neighbours were defined as

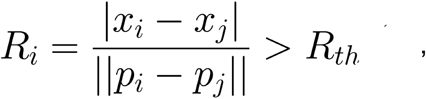

where *p_i_* and *p_j_* are nearest neighbours embedded in *m* dimensions, *x_i_* and *x_j_* are the same points embedded in *m* + 1 dimensions and *R_th_* is the threshold (*R_th_* = 10). The embedding dimension was defined for each dataset as the *m* at which the fraction of false nearest neighbours approached 0.

We then approximated the largest Lyapunov exponent (*λ*), which estimates the divergence of nearby trajectories in phase space, a defining feature of chaos (Babloyantz & Destexhe, 1986; Rosenstein et al., 1993). We firstly locate the nearest neighbour for each point along the reconstructed attractor expressed as

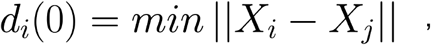

where *d_i_*(0) is the initial distance between X*_i_* and its nearest neighbour X*_j_*. From this process we calculate the change in distance over time for all nearest neighbours along the attractor which for each *t* is

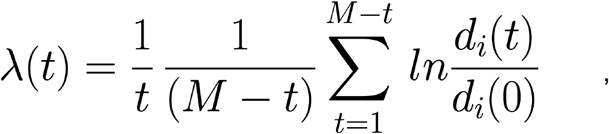

where *M* is the number of timesteps of the embedded attractor (Rosenstein et al., 1993; Sato et al., 1987). The largest Lyapunov exponent is thus the mean separation rate for nearest neighbour points over a given time course.

Given that we have full control of our *in silico* system, *λ* can be calculated in our model by perturbing the network and following trajectories over time, estimated as

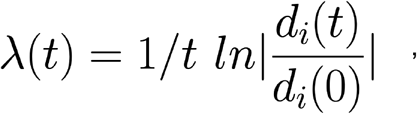

where *t* is the number of timesteps after the perturbation was applied.

#### Eigenspectrum analysis

The eigenspectrum function captures the variance of the n^th^ principal component, which is calculated by performing principal components analysis on a *k x m* matrix of *k* neurons and *m* time frames. Eigenspectra were calculated on raw fluorescence traces. To estimate the speed of the underlying dynamics, we define the velocity as the normalised Euclidean distance travelled per unit time across the whole neuronal population.

In order to study the relationship between eigenspectra and the velocity of the dynamics we use a 1-dimensional model introduced by Stringer et al. (2019). We define a function *f*(*x*) which describes the variance of *n* components across *x* samples as

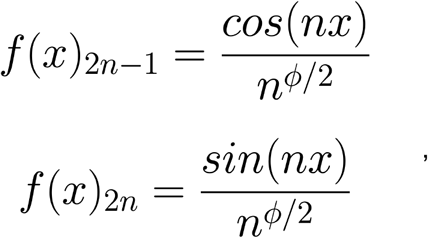

giving rise to covariance eigenvalues that follow n^*-ϕ*^. The quantity *n* was uniformly distributed between 0 and 2π. Changing *ϕ* changes the degree of variance captured by earlier versus later components, and thus alters the slope of the eigenspectrum. In order to visualise the effect of increasing *ϕ* on the velocity of the dynamics, we randomly project data matrix *X* (*n* x *x*) through a 3 dimensional randomised weight matrix *W* (3 x *n*).

#### Metastability analysis

In order to assess the number of metastable states we adapt methods used in a previous study (Haldeman & Beggs, 2005). A metastable state is defined as a set of state vectors that are more similar to one another than expected in a random system. To identify metastable states in our empirical data we performed affinity propagation across all state vectors for each condition. Affinity propagation is a clustering method which uses message passing to identify cluster exemplars, and which does not require cluster number to be defined *a priori*. Any clusters for which the number of state vectors belonging to that cluster was equal to 1 were removed. To calculate the similarity between state vectors that belong to a cluster we use

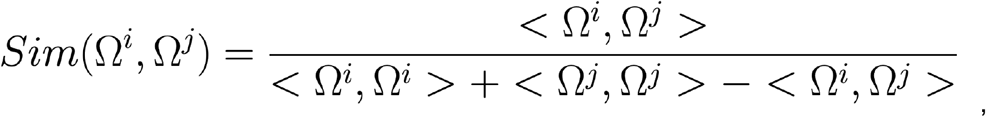

where Ω is a state vector and <, > denotes the dot product. For clusters to be identified as metastable, they needed to show greater similarity than expected by chance. We estimated the chance level similarity by performing affinity propagation clustering and similarity calculation on a null network using event-count matched shuffling. Any cluster with higher average similarity than the null network was declared a metastable state. One dataset failed to return any clusters for the *spontaneous* block and therefore we removed all conditions relating to this fish for further metastability analyses. To assess the mean dwell time in each metastable state we calculated the mean number of time frames in which the brain resided in a given state. In order to estimate the dwell time expected by chance for a system with *n* clusters, we randomly drew values between 1 and *n* from a uniform distribution, *t* times. The *δ* dwell time for each dataset was calculated as the empirical dwell time minus the null dwell time.

#### Statistical Tests and Software

D’agostino’s K^2^ test was used to test for normality in data distributions (α = 0.05). Paired t-tests or Wilcoxon signed-rank tests were used to compare *spontaneous, focal* ictal, *generalised* ictal, state transition, and human EEG datasets in cases of normality, and non-normality respectively (α = 0.05). Independent t-tests or Mann-Whitney U tests were used to compare different network models in cases of normality, and non-normality respectively (α = 0.05). Bonferroni corrections were used to control for false positives due to multiple comparisons.

In order to compare empirical and null avalanche distributions, we calculated the Kolmogorov Smirnov (KS) distance of each empirical distribution to the mean of its 50 null distributions, with the mean KS distance of each null against its own mean. To compare network response properties across different models we generated 50 simulations for each parameterised model, comparing the means of each simulation across model conditions. To calculate the effect of changing the network topology parameter *m* on network response properties, we parameterised the network to *preictal* levels before increasing *m* in steps to the levels of the *network topology* model, while keeping other parameters fixed. We performed 50 simulations for each *m*. We calculated the correlation between *m* and network response properties using Pearson’s correlation coefficient.

Data was analysed using custom code written in Python. Image registration and cell segmentation was performed using suite2p (Pachitariu et al., 2017). Neural network simulations were run using Brian2 (Stimberg et al., 2019). Statistical hypothesis tests were performed using scipy. Graphs were generated using matplotlib and seaborn.

## Supplementary Information

**S1.**
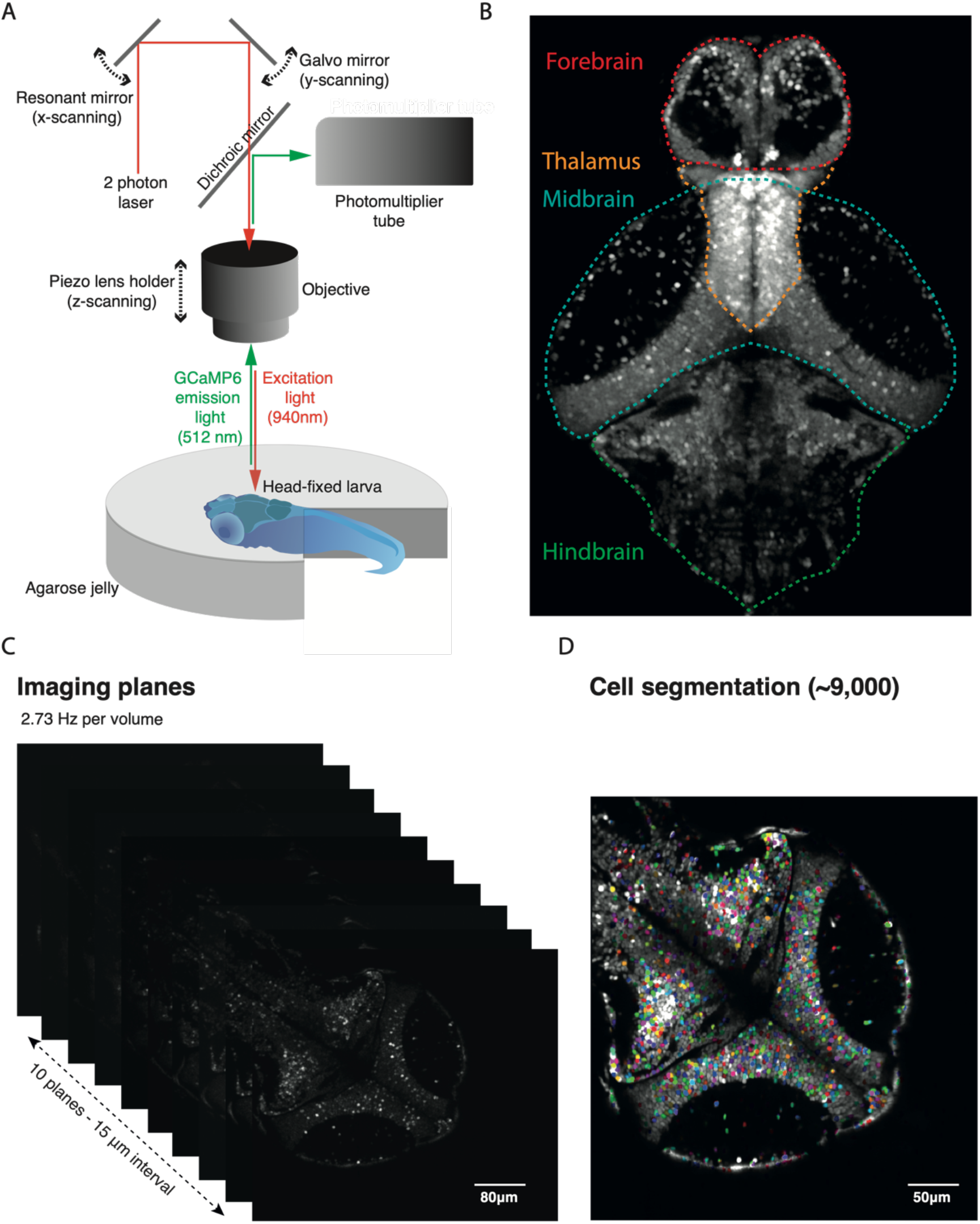
2 photon imaging setup and data processing. (A) 2 photon imaging setup with head-fixed larval zebrafish. (B) Max projection across zebrafish volume taken with 2 photon microscope, demonstrating coverage of major brain regions. (C) Imaging was captured across 10 planes with 15 um spacing at an imaging rate of 2.73 Hz per volume. (D) Stacks from each plane were rigid registered together and single cell segmentation was performed using suite2p.

**S2.**
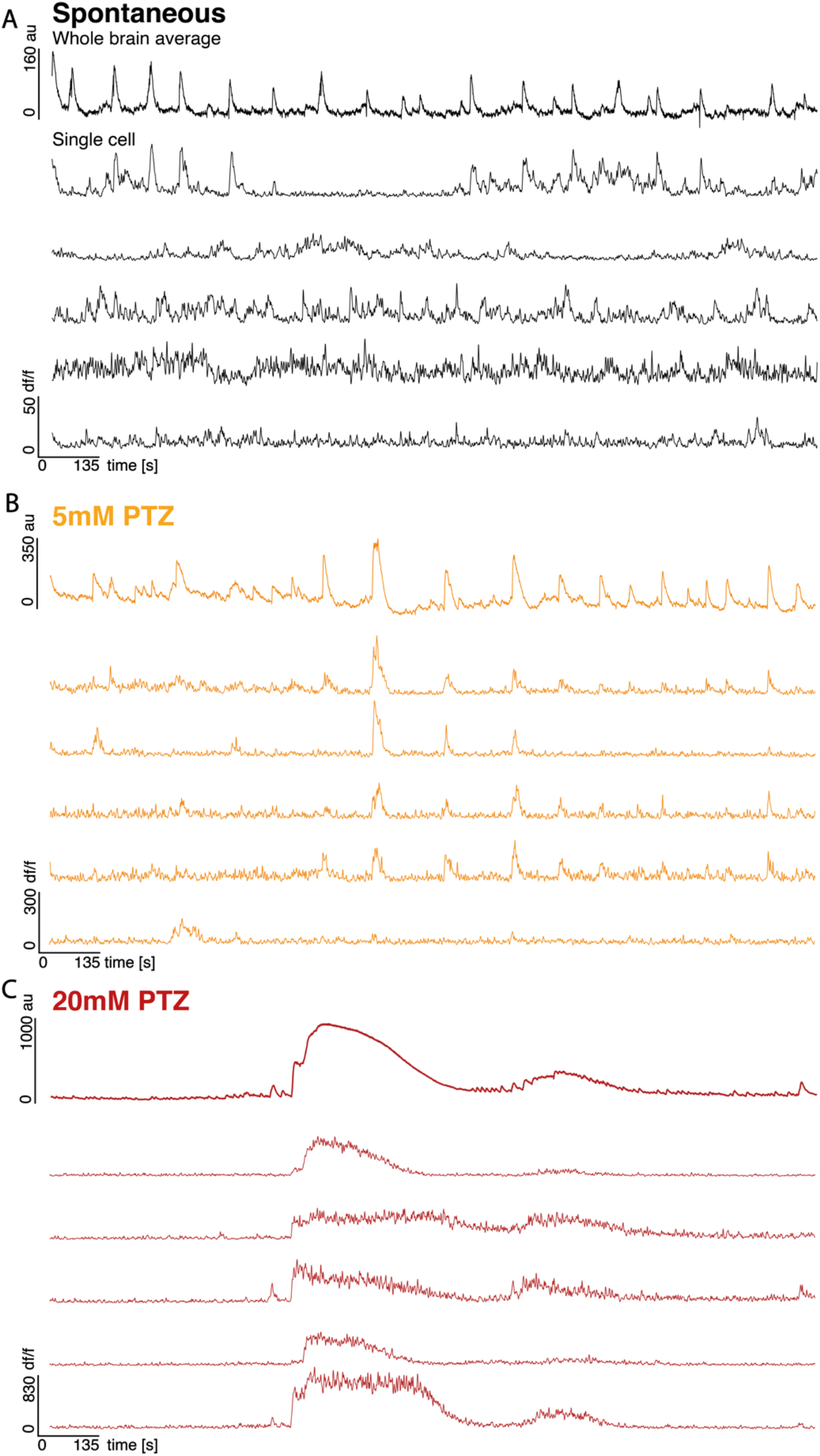
Single cell and whole brain dynamics of EI balance and imbalance. (A-C) Whole brain average (top trace) calcium fluorescence and representative single cell traces (bottom 4 traces) for *spontaneous* (black), *5mM PTZ* (orange) and *20mM PTZ* (red) conditions for the entire 30 minute recording period for a representative fish. EI balanced dynamics (A) showed low correlation interspersed with brief high correlation events. EI imbalanced dynamics showed regular bursts of correlation (B) and sustained, high amplitude synchronous cascades (C).

**S3.**
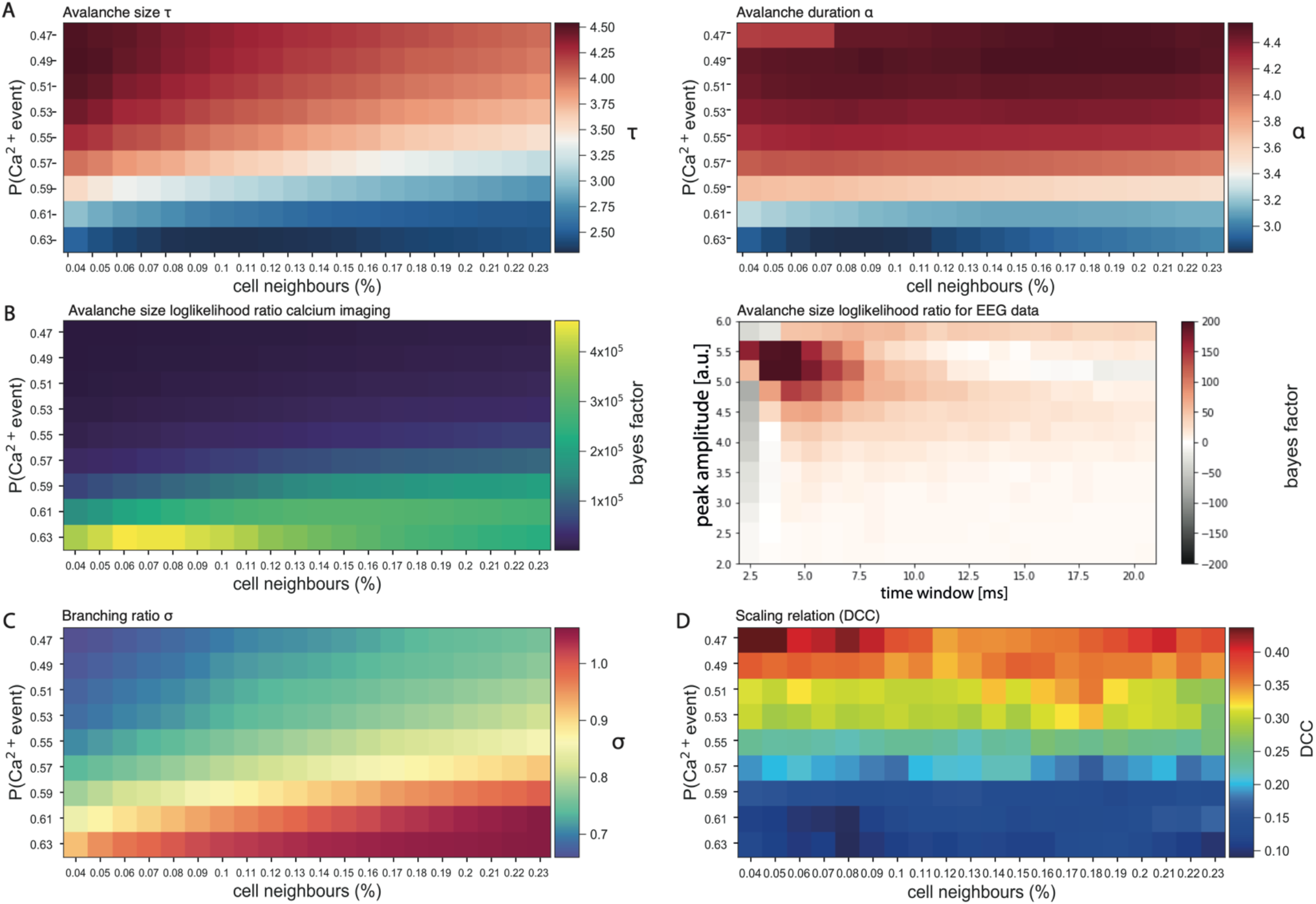
Critical statistics are parameter dependent. Critical statistics vary across different model parameter combinations - the event probability in the Hidden Markov Model (P(Ca2+ event)) and the % of closest neurons assigned to a neighbourhood. (A) Exponents for avalanche size (left) and duration (right) vary according to both event probability and neighbourhood size. (B) Avalanche size distributions for zebrafish calcium imaging (left) and human EEG (right) data are better explained by power law than by lognormal forms across a majority of parameters as shown by positive Bayes factor for majority of parameter combinations. Bayes factors are calculated as loglikelihood ratios for power law vs lognormal distributions. (C) Branching ratio varies according to both event probability and neighbourhood size. (D) Exponent relation as measured by the deviation from criticality coefficient (DCC) varies according to event probability.

**S4.**
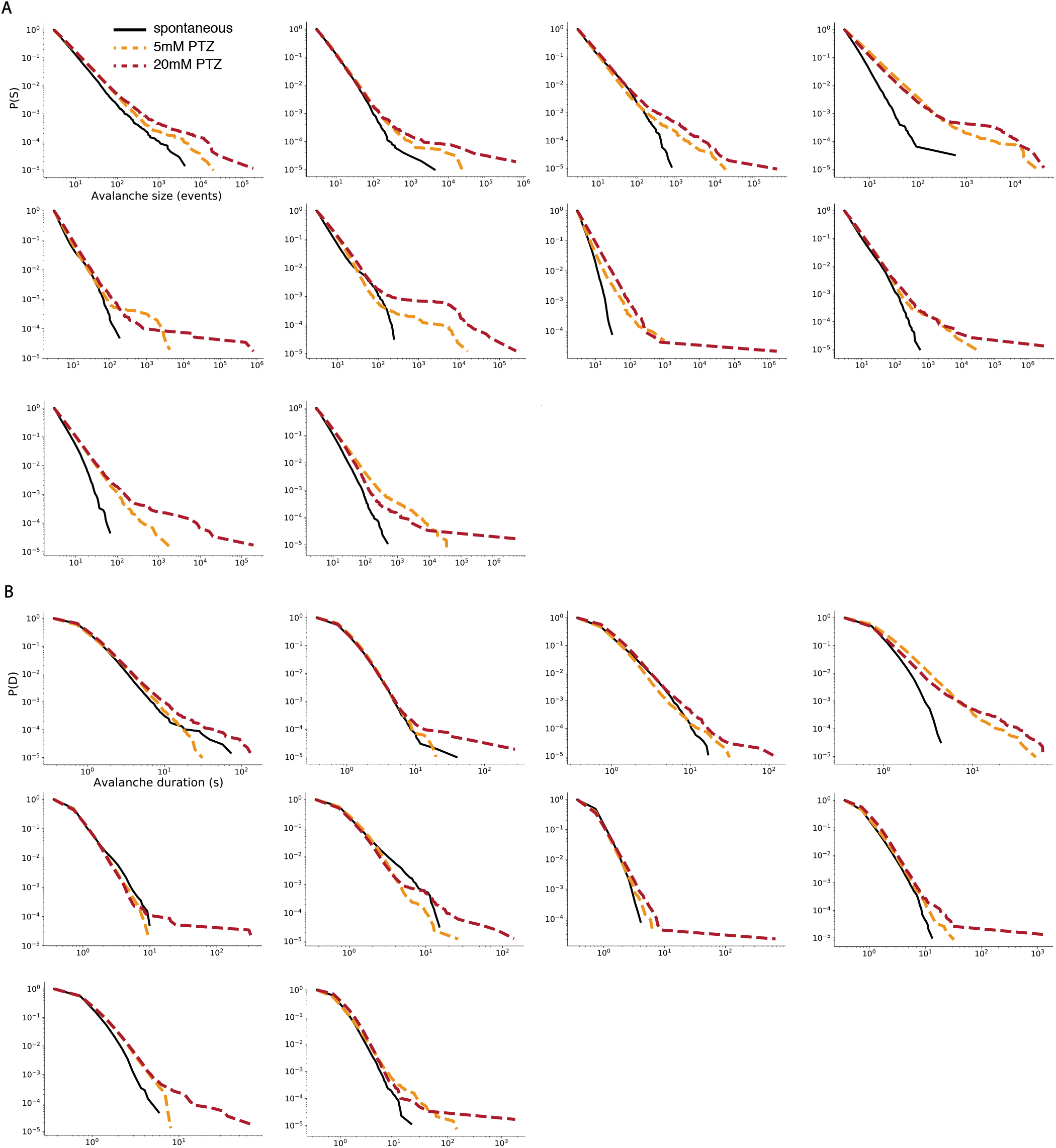
Avalanche distributions increase in slope during EI imbalance. (A-B) Complementary cumulative distribution functions for *spontaneous* (black, solid line), *5mM PTZ* (orange, dotted line), and *20mM PTZ* (red, solid line) periods for avalanche size (A) and duration (B) for each fish.

**S5.**
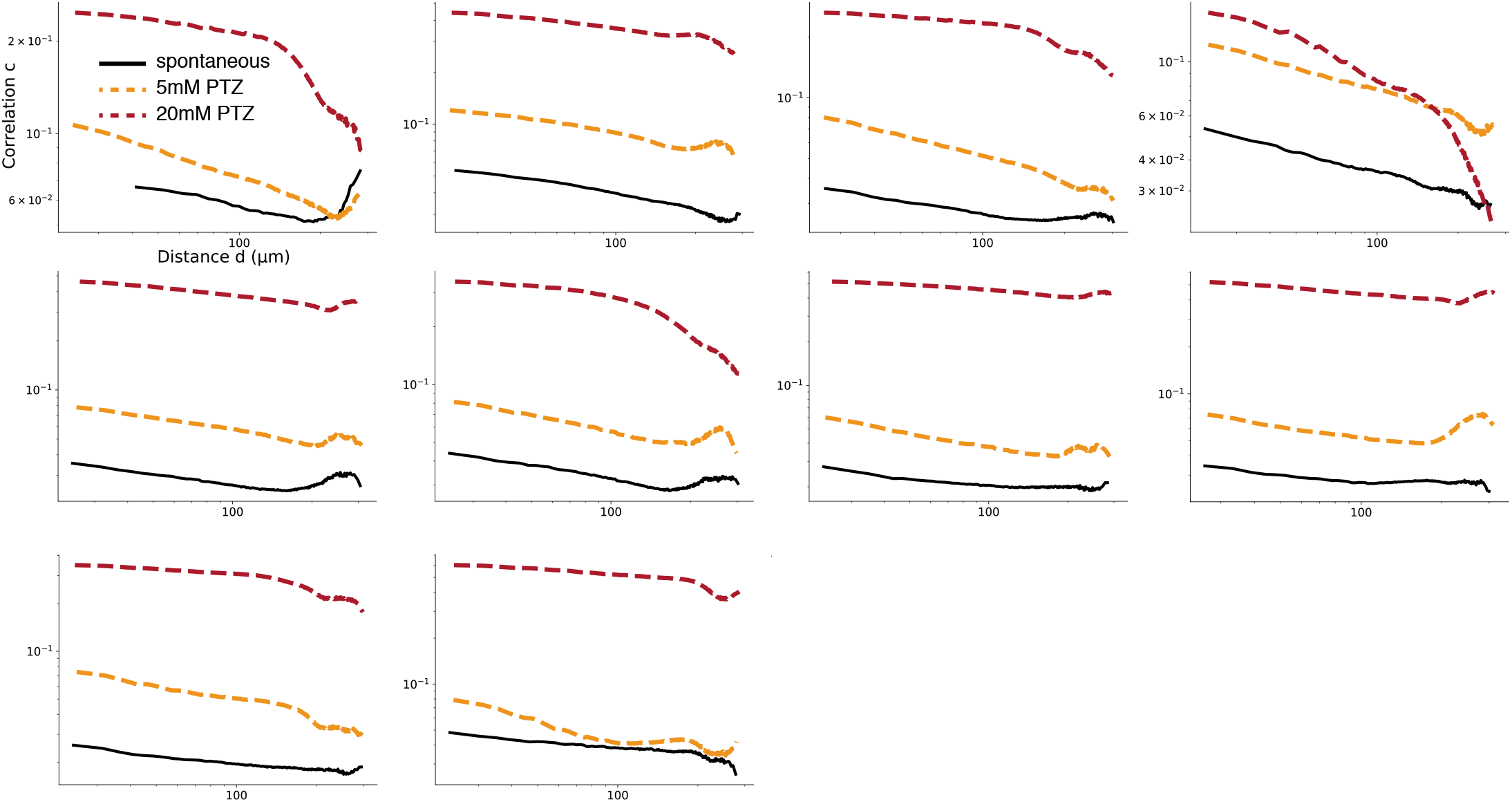
Loss of correlation function power-law relationships due to EI imbalance. Correlation functions for *spontaneous* (black, solid line), *5mM PTZ* (orange, dotted line) and *20mM PTZ* (red, dotted line) periods for each fish. *5mM PTZ* and *20mM PTZ* datasets are less well fit to power-laws than spontaneous datasets.

**S6.**
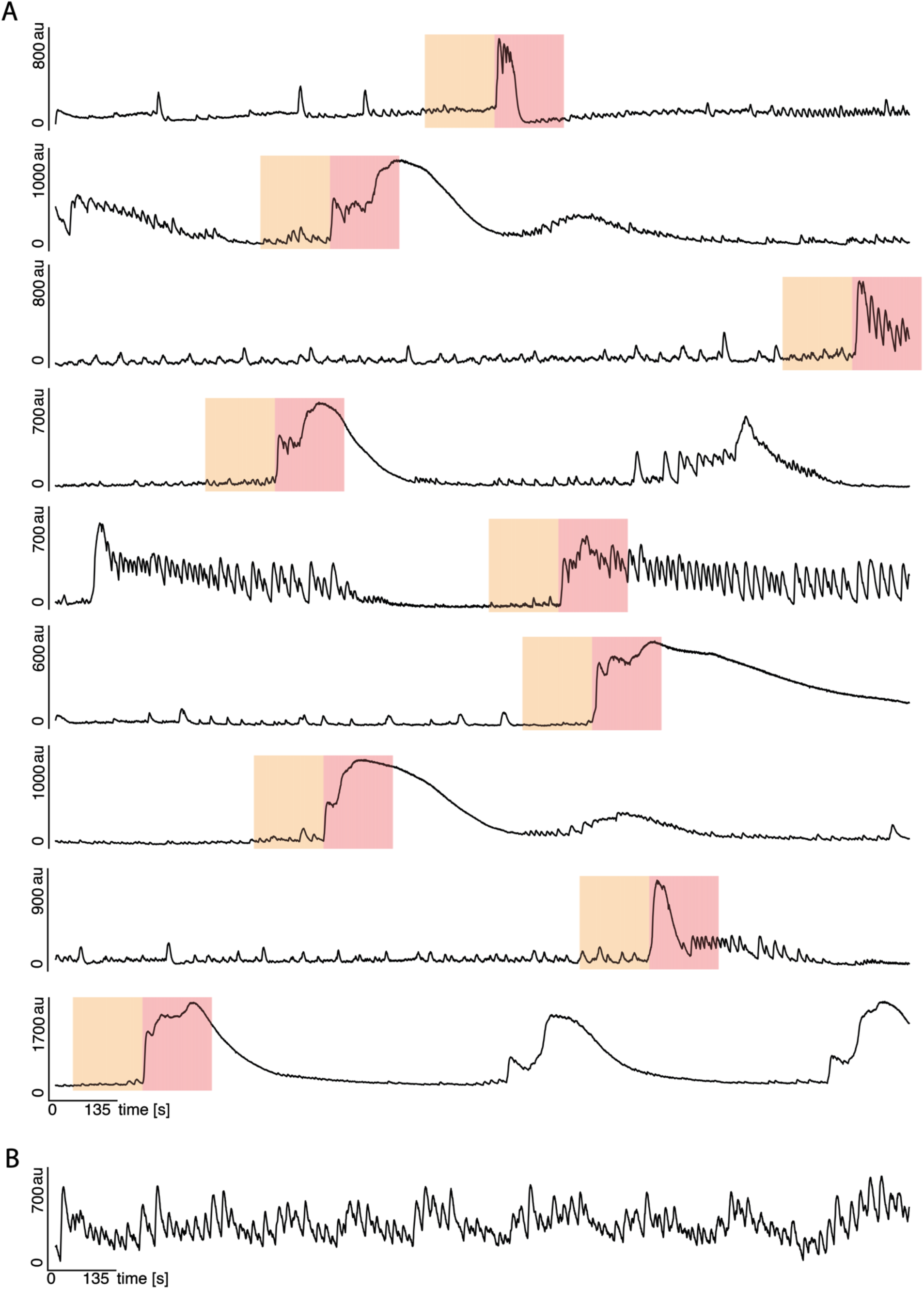
Identification of seizure state transitions. (A) Mean whole brain traces showing 400 frame *pre-ictal* (orange) and *ictal-onset* (red) periods used for 9 datasets. Start of state transition was identified using a 30 frame sliding window, with the window with the maximal difference over the preceding 29 frames identified as the start of the *ictal-onset* period. (B) One dataset without clear state transitions from *pre-ictal* to *ictal-onset* state was removed.

